# The neurodevelopmental disorder-linked PHF14 complex that forms biomolecular condensates detects DNA damage and promotes repair

**DOI:** 10.1101/2021.10.12.462922

**Authors:** Eva-Lotta Käsper, In-Young Hwang, Helga Grötsch, Herman Kar Ho Fung, Aurélien A. Sérandour, Niccolò Arecco, Ronald Oellers, Patryk Poliński, Almudena Garcia Gomez-Monedero, Christoph W. Müller, Kyung-Min Noh

## Abstract

Numerous chromatin-associated proteins have been linked to neurodevelopmental disorders, yet their molecular functions often remain elusive. PHF14, HMG20A, TCF20 and RAI1 are components of a putative chromatin-associated complex and have been implicated in neurological disorders. Here, we found that Phf14 knockout embryonic stem cells and neural progenitor cells exhibit impaired cell cycle progression and proliferation, inadequate protection of stalled replication forks, and decreased DNA repair. The PHF14 complex rapidly assembles at DNA damage sites and binds to DNA through HMG20A. The PHF14 complex forms DNA-containing phase separated droplets in vitro, where TCF20 facilitates droplet formation. Furthermore, TCF20 maintenance at DNA damage sites is destabilized upon pathological mutation. Our results suggest that the PHF14 complex contributes to DNA damage repair by sensing damaged sites and forming biomolecular condensates, thus supporting cell cycle progression, especially in neural progenitor cells whose spatiotemporal pool is critical for proper brain development.

## Main

Neurodevelopmental disorders arise from genetic and/or environmental insults that affect brain development and function. Mutations associated with neurodevelopmental disorders are often in genes within common pathways, such as chromatin remodelers and DNA repair. Members of a putative chromatin regulatory complex comprising PHF14, HMG20A (iBRAF), TCF20 (SPBP) and RAI1 (hereafter PHF14 complex),^1,2^ are strongly linked to neurodevelopmental disorders. *PHF14* encodes the PHD finger protein 14 and resides at the 7p21.3 locus, which is deleted or duplicated in some cases of Dandy-Walker syndrome, a congenital brain malformation involving the cerebellum^3^. Missense, nonsense and deletion mutations in *TCF20*, which encodes transcription factor 20, were identified in intellectual disability and autism spectrum disorders^4–7^. Retinoic acid induced 1 (RAI1) is a paralog of TCF20, and mutations and deletions in *RAI1* cause Smith-Magenis syndrome, whereas *RAI1* duplications cause Potocki-Lupski syndrome^8^. Both are characterized by mild to moderate intellectual disability, disrupted sleep patterns, infantile hypotonia, obesity and distinctive facial features^9^.

The PHF14 complex was identified in HeLa cells, but its physiological regulation and function are not known^1,2^. TCF20 is highly expressed in the brain and is localized to the nucleus^10^. It is a putative transcriptional coregulator, yet its exact role within the nucleus is unclear. RAI1 has been implicated in the transcriptional regulation of a small subset of neuronal genes^11,12^. Given that RAI1 and TCF20 are both components of the PHF14 complex, the similar developmental disorders caused by their mutations might arise due to dysfunction of the complex. The function of PHF14 is unknown, but it is deleted in biliary tract cancer cell line OZ^13^, biallelically inactivated in colon cancer cell lines^14^ and its higher expression levels in lung cancer patients correlate with poor survival^15^. HMG20A has been associated with the LSD1-CoREST complex, in which its paralog HMG20B is reported to be a subunit, and found to regulate genes required for neuronal differentiation^16^.

Each of the PHF14 complex components have putative DNA or histone interacting domains, suggesting its association with chromatin. HMG20A contains a high mobility group (HMG) box domain and the HMG-box in its paralog HMG20B binds four-way junction (4WJ) DNA without sequence-specificity^17^. PHF14 contains several PHD fingers: a PHD finger - Zn knuckle - PHD finger (PZP) domain and other zinc fingers at its C-terminus. Similar PZP domains have been shown to recognize H3 tails^18^. TCF20 and RAI1 contain an extended PHD (ePHD), also called ADD domain at the C-terminus. ADD domains of DNMT3A and ATRX were shown to bind modified H3 tails^19,20^.

Here, we set out to understand the function of this minimally characterized protein complex (PHF14, HMG20A, TCF20, RAI1), to decipher how mutations and copy number variations of the corresponding genes lead to various overlapping neurodevelopmental disorders. Using genomic, proteomic, and biochemical approaches, we discovered that the PHF14 complex regulates proliferation of neural progenitor cells. Mechanistically, we found that this complex phase separates, binds damaged DNA, and facilitates DNA repair.

## Results

### Distinct Phf14 complexes form in ESCs and NPCs

To investigate the composition of the PHF14 complex during development, we examined mouse embryonic stem cells (ESCs) and neural progenitor cells (NPCs) derived from ESCs with retinoic acid treatment^21^. We performed immunoprecipitation followed by quantitative mass spectrometry analysis (IP-MS) using an antibody against Phf14. As a control, we generated *Phf14* knockout (KO) ESCs using CRISPR-Cas9, which removed exon 5 of the *Phf14* gene encoding a part of the structured PZP domain, resulting in complete loss of the full-length protein due to frameshift-induced nonsense-mediated decay (Extended Data Fig. 1a). Enrichment analysis (Phf14 IP-MS in wild-type vs. *Phf14* KO) revealed that Phf14 interacted with Hmg20a, Tcf20, and Rai1 in both ESCs and NPCs (Fig. 1a). While HMG20A has been proposed to replace its homolog HMG20B in the LSD1-CoREST complex^22^, we did not observe interactions between Phf14 and LSD1-CoREST complex members in either ESCs or NPCs. However, we found that Tcf20 expression and binding to Phf14 were reduced in NPCs relative to ESCs (Fig. 1b and Extended Data Fig.1b, c). In contrast, Rai1 expression and binding to Phf14 were increased in NPCs relative to ESCs (Fig. 1b and Extended Data 1b, c). Overall, these data suggest that the composition of the Phf14 complex changes during neurodevelopment.

**Fig. 1:**
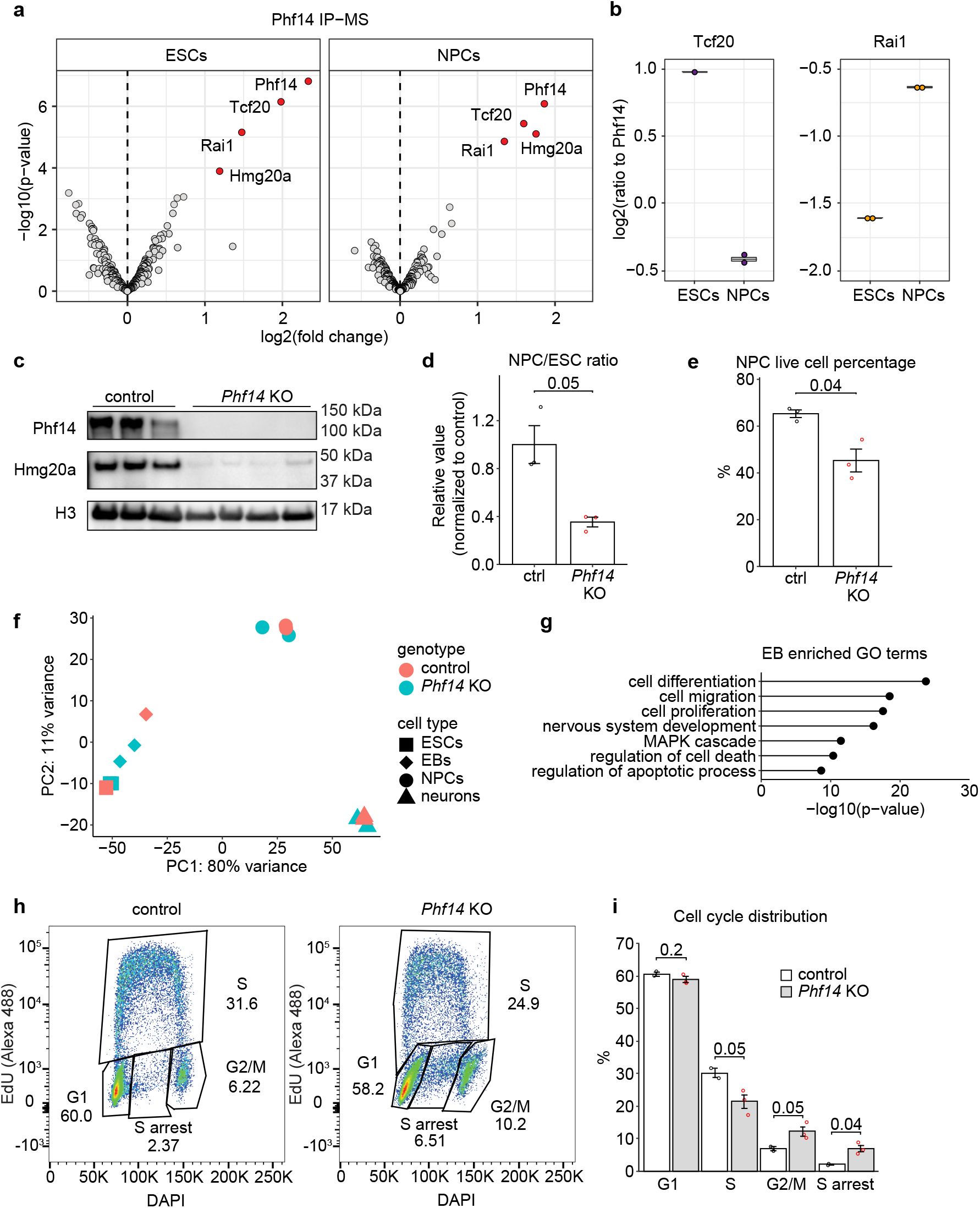
Phf14 forms a complex with Rai1, Tcf20 and Hmg20a, and *Phf14* KO NPCs exhibit cell proliferation defects and increased cell cycle arrest. **a**, Phf14 interacting proteins in ESCs and NPCs identified by quantitative tandem mass tag (TMT) spectrometry. Immunoprecipitation samples from control cells are compared to *Phf14* KO. Red - significantly enriched proteins with adj. p-value < 0.05, grey – not significant. P-values and fold changes were obtained from limma statistical analysis. n = 2. **b**, Tcf20 and Rai1 normalized to Phf14 signal (with the same TMT-label) in ESCs and NPCs, related to (**a**). **c**, Western blot of nuclear lysate of control and *Phf14* KO cells. The experiment was repeated three times with similar results. Histone H3 serves as a nuclear loading control. **d**, Control-normalized values of number of NPCs derived from the same amount of ESCs at the start of differentiation. **e**, Trypan-blue based live-cell percentage of NPCs. n = 3, two-sided unpaired Student’s t-test. **f**, Principal component analysis plot of *Phf14* KO and control samples including all timepoints. (day 0 – ESCs, day 4 – embryoid bodies (EBs), day 8 – NPCs, day 12 – neurons). **g**, GO enrichment analysis for genes differentially expressed in *Phf14* KOs compared to controls on day 4. **h**, Representative cell cycle distribution plots for monolayer-differentiated NPCs (day 4). **i**, Quantified data related to (**h**). Two-sided unpaired student’s t-test was used for calculating p-values.

To determine if Rai1 interacts with Phf14 in NPCs, we introduced a HA-FLAG tag at the C-terminus of endogenous Rai1 (Extended Data Fig. 2a) to bypass the lack of a reliable antibody against Rai1 and performed IP-MS. Indeed, we identified Phf14 as a significant hit. The RAI1 C-terminal ePHD/ADD domain is occasionally lost in patients with Smith-Magenis syndrome^9^. We tested if the RAI1-ePHD domain interacts with a specific region of the Phf14 and found that the PZP domain of PHF14 was required for the interaction (Extended Data Fig. 2b). Although the RAI1-ePHD has been proposed to bind nucleosomes in a histone-tail-dependent manner^23^, we did not detect an interaction between RAI1-ePHD and any histone tails (Extended Data Fig. 2c), consistent with another report^11^.

We also examined whether the Rai1-ePHD is required to regulate gene expression. To do so, we used CRISPR-Cas9 to remove exon 4, which encodes the zinc finger in ePHD from the *Rai1* gene (Extended Data Fig. 2d), resulting in a truncated protein. Given that Smith-Magenis syndrome is a neurodevelopmental disorder, we differentiated *Rai1* exon 4 KO ESCs into glutamatergic neurons via NPCs^21^, and performed mRNA-seq, but observed minimal changes (17 genes down- and 14 up-regulated in neurons, none in NPCs) (Extended Data Fig. 2e). We infer that the RAI1-Phf14 interaction does not regulate gene expression in NPCs or neurons and might function in another cellular pathway.

### *Phf14* KO cells exhibit an altered cell cycle and increased S-phase arrest

To decipher the function of the Phf14 complex, we examined the *Phf14* KO ESCs. When we checked the Phf14 protein expression by immunoblot, in addition to the loss of Phf14, we observed a significant reduction in Hmg20a levels (Fig. 1c). This is likely due to protein degradation in the absence of Phf14, since mRNA levels of *Hmg20a* were only mildly reduced in *Phf14* KOs (Extended Data Fig. 1d). Thus, these cells have lost more than one protein in the Phf14 complex, disrupting the formation of the complex. We differentiated the *Phf14* KO and wild-type control ESCs into neurons, and collected cells at distinct timepoints (day 0 ESCs, day 4 embryoid bodies (EB), day 8 NPCs, day 12 neurons). Compared to controls, *Phf14* KO ESCs generated a lower yield of NPCs (Fig. 1d) and *Phf14* KO NPCs showed increased cell death (Fig. 1e). Yet, *Phf14* KO NPCs could differentiate into neurons (Extended Data Fig. 1e).

We performed mRNA-seq to evaluate transcriptome changes during neurodevelopment, and principal component analysis showed separation between the timepoints, as expected. Notably, the *Phf14* KO EBs clustered close to the ESCs (Fig. 1f), suggesting that they are slower to exit from pluripotency than controls. GO analysis of the differentially expressed genes (DEGs) in *Phf14* KO versus control EBs and ESCs revealed terms related to cell differentiation, migration, proliferation as well as nervous system development, regulation of cell death and apoptosis (Fig. 1g and Extended Data Fig. 1f).

We observed and validated the upregulation of the cell cycle regulators *Cdkn2a* and *Cdkn2b* upon *Phf14* KO (Extended Data Fig. 1g); these genes were also upregulated upon *PHF14* knockdown in a cancer cell line^24^. *Cdkn2a* encodes p16 and p19ARF and *Cdkn2b* encodes p15-INK4b, which promote cell cycle arrest, senescence or apoptosis^25^. To determine if *Phf14* KO affects the cell cycle, which might explain the lower yield of NPCs, we performed an EdU-incorporation assay. This assay must be performed on a monolayer of cells to ensure even uptake of EdU, so we differentiated the ESCs to neural progenitors following a monolayer differentiation protocol. Compared to the controls, the *Phf14* KO NPCs displayed an increased proportion of cells in G2/M phase and a reduced proportion of cells in S-phase. In addition, *Phf14* KO NPCs displayed an increased frequency of S-phase arrest when compared to wild-type NPCs (Fig. 1h, i), suggesting a perturbed cell cycle.

### Phf14 promotes DNA repair

We considered that the replication arrest, altered cell cycle, and proliferation defects of *Phf14* KO arise from issues with DNA damage checkpoints and repair. Indeed, PHF14 has been identified at stalled replication forks^26^, and, similar to known DNA damage response (DDR) factors, all members of the PHF14 complex undergo changes in their post-translational modifications (PTMs) in response to genotoxic agents, including increased poly ADP-ribosylation (PARylation), decreased small ubiquitin-like modifier (SUMOylation) and changes in phosphorylation (Supplementary Table 1),^27–33^.

To determine if Phf14 protects cells from replication stress, we treated cells with the DNA replication inhibitor hydroxyurea (HU) and performed a DNA fiber spreading assay^34,35^. HU treatment stalls replication forks. Briefly, we sequentially labeled replicating DNA with IdU followed by CldU, treated cells with HU, and examined the CldU/IdU ratio before and after treatment. The CldU/IdU ratio correlates with the stability of the stalled replication forks. Before HU treatment, *Phf14* KO and wild-type ESCs and NPCs displayed similar CldU/IdU ratios. However, after HU treatment, the CldU/IdU ratio was lower for *Phf14* KO cells compared to controls, particularly in NPCs (Fig. 2a). These data suggest that Phf14 stabilizes stalled replication forks in ESCs and NPCs.

**Figure 2.**
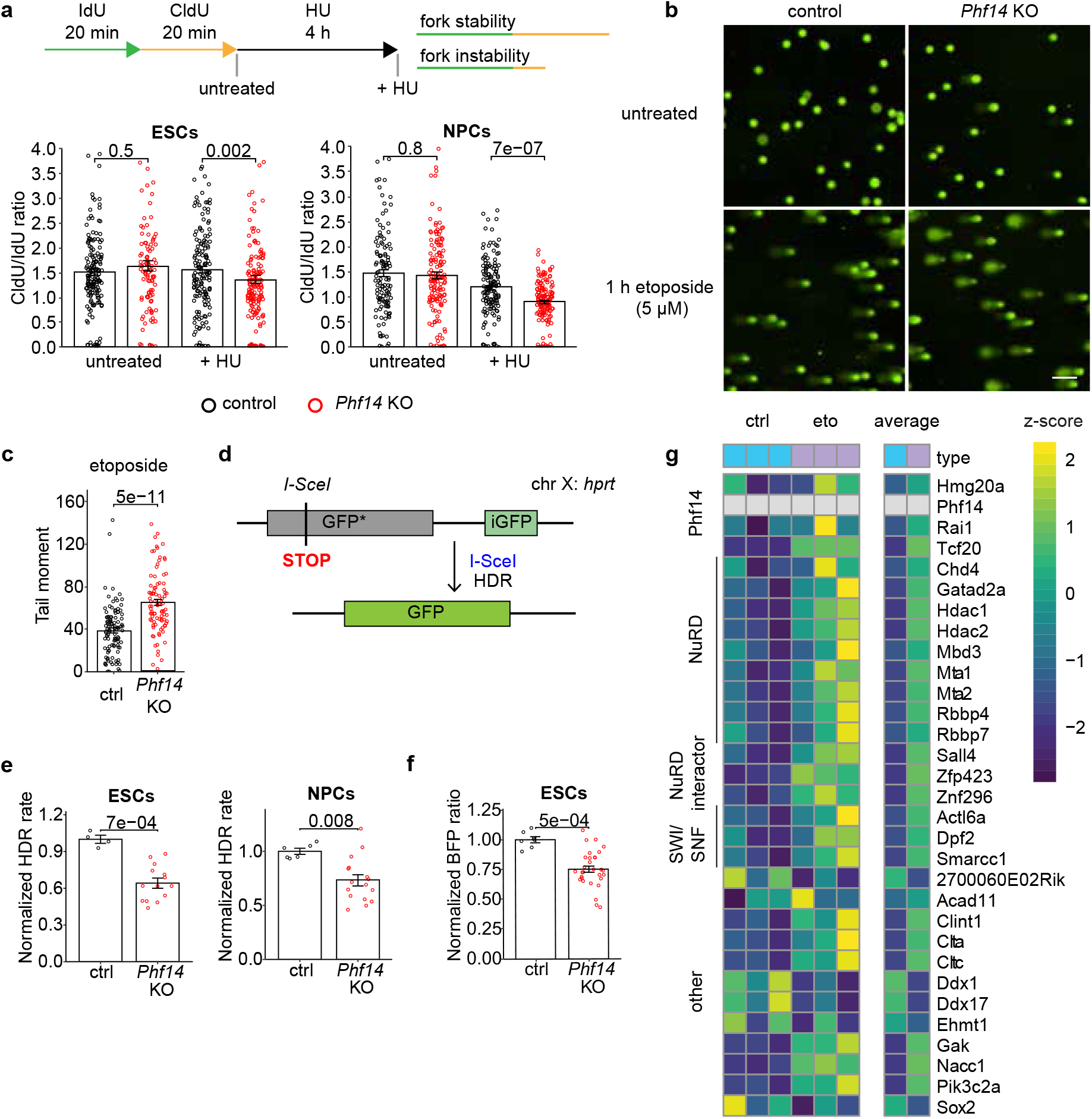
*Phf14* KO cells have defects in DNA damage repair and impaired stalled replication fork protection. **a**, CldU/IdU track ratios from DNA fiber spreading using *Phf14* KO and control ESCs and monolayer-differentiated NPCs. Over 100 tracks were quantified per condition. Two-sided Wilcoxon rank-sum test was used for calculating p-values. **b**, Widefield microscope images of an alkaline comet assay of control and *Phf14* KO neurons. Cell were collected either untreated or 1 h after 5 μM etoposide treatment. Scale bar = 100 μm. **c**, Comet tail moments after etoposide treatment, quantified using CometScore 2.0 software. Vast majority of untreated comets had a tail moment of 0. **d**, Principle of hprtDRGFP reporter lines. The *GFP* sequence contains a stop codon that disrupts the expression of GFP near an I-SceI restriction site. The incomplete iGFP can be used as a repair template in HDR to restore a functional GFP. **e**, Homology-directed repair rates in *Phf14* KO and control hprtDRGFP ESCs and monolayer-differentiated NPCs after double strand break induction using lentivirus-transduced I-SceI-T2A-BFP enzyme. GFP-positive cells were counted using flow cytometry and the HDR rate was calculated as the ratio of GFP-positive cells among cells gated for BFP for successful transduction. **f**, Amount of BFP-positive cells (cells that have I-SceI expression) in the measured population in control and *Phf14* KO cells. Same amount of cells was plated the day before transduction with the same amount of lentivirus. **g**, Heatmap showing the z-scores from bait ratios (identified protein signal to Phf14 signal in the sample) of proteins that were significantly enriched in both untreated and etoposide treated IPs over KO controls. Phf14 interacting proteins were identified in ESCs without and after 10 μM etoposide treatment for 1 h identified by quantitative tandem mass tag (TMT) spectrometry. n = 3 controls, 2 KOs.

To determine if Phf14 promotes genomic stability outside of S-phase, we performed a comet assay in postmitotic neurons. Cells were treated with the DNA-damaging agent etoposide and DNA fragmentation levels were determined by single-cell electrophoresis and quantification of the comet tail moment. In the undamaged condition, all but a few cells had a comet tail moment of 0 for both control and *Phf14* KO neurons. The tail moment increased to 64.9 for *Phf14* KO neurons but only to 38.8 for wild-type neurons (Fig 2b, c), indicating that Phf14 also protects post-mitotic cells from DNA damage. A similar but milder trend was also observed in the *Rai1* ex 4 KO neurons with a post-etoposide tail moment of 46.7 (Extended Data Fig. 2f).

To further determine if double-strand break repair was impaired in the *Phf14* KO cells, we generated lines expressing the homology directed repair reporter DR-GFP. This reporter contains an inactive GFP gene due to a stop codon next to an I-SceI site and a repair template that can be used by the cells to restore a functional GFP sequence^36^ (Fig. 2d). We introduced the I-SceI enzyme and BFP as a transduction marker to cells using lentivirus, and measured the GFP:BFP ratio by flow cytometry to detect successful homology-directed repair among the cells that had a double-strand break. Compared to controls, *Phf14* KO cells showed a decreased rate of homology-directed repair in both ESCs and NPCs (Fig. 2e). Additionally, the BFP-positive *Phf14* KO ESCs exhibited decreased proliferation compared to BFP-positive wild-type ESCs, suggesting slower DNA repair and resumption of cell division and/or increased cell death with I-SceI induced double-strand breaks (Fig. 2f). Thus, the 25-30% decrease in homology-directed repair (Fig. 2e) might underestimate the true defect.

Given the observation that Phf14 promotes genome stability, we determined if DNA damage affects the Phf14 interactome. Specifically, we used the Phf14 antibody for IP-MS of control and etoposide-treated ESCs. Etoposide treatment led to new or increased protein interactions of Phf14, particularly with members of the NuRD and SWI/SNF chromatin remodeling complexes, and proteins that interact with those remodelers (Fig. 2g). Phf14 was previously reported to interact with NuRD^1^ but not with SWI/SNF. We also noted that Phf14 showed an increased association with Hmg20a, Rai1 and particularly Tcf20 after etoposide treatment. Together, these data suggest that the Phf14 complex is stabilized upon DNA damage and promotes DNA repair.

### The PHF14 complex components rapidly localize to DNA damage sites

Remodeler complexes are rapidly recruited to DNA damage sites^37^. To investigate whether the Phf14 complex is recruited to damaged loci, we used UV-laser micro-irradiation to locally induce DNA damage in monolayer-differentiated NPCs. We detected Hmg20a and Phf14 with available antibodies, and the endogenously tagged HA-Rai1 with anti-HA, but there are no reliable antibodies available for Tcf20. We found that all three endogenous proteins were enriched at the laser-induced DNA damage track marked by gammaH2A.X (Fig. 3a).

**Fig. 3:**
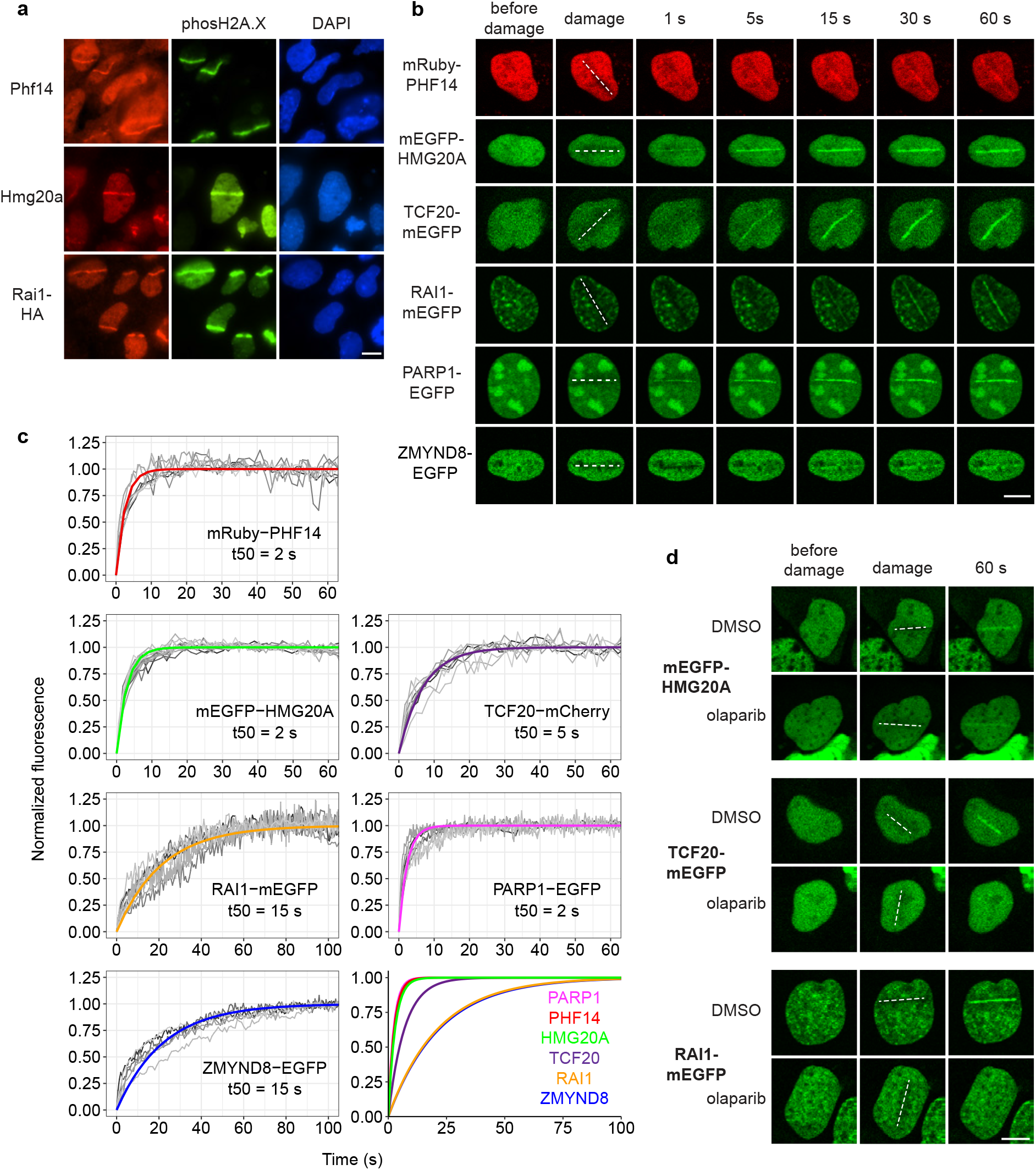
PHF14 and HMG20A localize to DNA damage sites simultaneously at a comparable speed to PARP1, followed by TCF20 and RAI1. **a**, Immunofluorescence staining in monolayer-differentiated NPCs after laser microirradiation. Anti-phospho-histone H2A.X (Ser139) signal serves as a DNA damage marker. Cells were fixed 5-15 minutes after damage generation. **b**, Live-cell imaging of fluorescently tagged PHF14 complex proteins and PARP1 and ZMYND8 as positive controls (transient transfection into U2OS cells). DNA damage was induced with 355 nm laser at the indicated location. **c**, Quantified recruitment kinetics from live-cell laser microirradiation experiments for the PHF14 complex and controls (in U2OS for RAI1, PARP1 and ZMYND8, other proteins from HEK293T cells, no difference was observed between different cell lines in terms of recruitment kinetics). Time is set to 0 at the time of DNA damage induction and intensities are normalized such that the maximum reached is 1. A first-order exponential equation f(t) = 1-exp(−t / τ) was fitted to all data per protein. The time required for the normalized fluorescence intensity to reach 50% of its maximal value is represented as t50. The last panel includes fitted curves from the other panels only. n = 8 (PHF14), 14 (HMG20A), 8 (TCF20), 8 (RAI1), 8 (PARP1), 6 (ZMYND8). **d**, Laser microirradiation in transiently transfected U2OS cells after treatment with 1 μM olaparib for 1 h or equivalent amount of DMSO. All scale bars = 10 μm.

To further investigate the recruitment of the Phf14 complex to damaged DNA in live cells, we transiently expressed tagged versions of the four proteins in cell lines. All members of the PHF14 complex showed fast recruitment to the DNA damage track induced by UV-laser micro-irradiation, as did the known DDR factors PARP1 and ZMYND8 in U2OS cells (Fig. 3b)^38,39^. The fluorescent proteins alone (e.g., mEGFP) did not accumulate at DNA damage sites, as expected (Extended Data Fig. 3c). PHF14 complex enrichment at DNA damage sites persisted for at least one hour after damage induction (Extended Data Fig 3a, b).

We found that HMG20A and PHF14 localized to damage sites at the same speed, reaching a half-maximum accumulation in approximately 2 seconds. TCF20 and RAI1, were slightly slower, reaching half-maxima in 5 and 15 s, respectively (Fig. 3c). The recruitment speed of PARP1 in our setup was similar to HMG20A and PHF14, with a half-maximum accumulation time (t_50_) of 2 s, in agreement with the literature^38^. ZMYND8, while still one of the faster responders to DNA damage, was recruited slower than the other proteins, except for RAI1, with a t_50_ of also 15 s, which is also consistent with previous reports^40^. These data indicate that the PHF14 protein complex is likely one of the first responders at DNA damage sites.

PHF14 and HMG20A localize to damage sites simultaneously, so we sought to determine if PHF14 was necessary for the recruitment of HMG20A by examining *Phf14* KO ESCs. We did not observe recruitment of endogenous Hmg20a in *Phf14* KO cells (Extended Data Fig. 3d), possibly because Hmg20a levels are reduced in these cells (Fig. 1c). Indeed, when we expressed mEGFP-HMG20A in *Phf14* KO ESCs, we found that it was recruited to DNA damage sites (Extended Data Fig. 3e, f). These data suggest that PHF14 is not absolutely required to recruit HMG20A to damage sites, but might be necessary for the stability and specificity of the complex with HMG20A.

PARylation is necessary for the recruitment of many DDR proteins to DNA damage sites^41,42^ and the PHF14 complex is PARylated in response to genotoxic stress (Supplementary Table 1). To determine if the recruitment of PHF14 complex components to DNA damage also requires PARylation, we treated cells with the PARP inhibitor olaparib before laser micro-irradiation. We found that HMG20A and PHF14 were still recruited to damage sites in the presence of olaparib (Fig. 3d, Extended Data Fig. 3g). However, RAI1 and TCF20 showed no or minimal recruitment in olaparib-treated cells (Fig. 3d), indicating that PARylation is important for their localization to damage sites. Taken together, the kinetics as well as the PARylation-sensitivity suggest that the PHF14 complex components localize to damage sites in a stepwise manner, with HMG20A and PHF14 arriving first.

### PHF14 and HMG20A form a binary complex that interacts with TCF20A or RAI1

PHF14 and HMG20A interact strongly with each other to form a binary core complex that rapidly localizes to DNA damage sites. To further characterize this core complex, we purified the proteins from insect cells (Extended Data Fig. 4a). PHF14 and HMG20A were present in the same elution fractions from both ion-exchange and size-exclusion chromatography, suggesting that they interact in solution. To identify the interacting regions, we cross-linked the purified proteins and performed MS, which revealed that the C-terminal portion of PHF14 interacts with the HMG-box and coiled coil of HMG20A (Fig. 4a, Extended Data Fig. 4b).

**Fig. 4:**
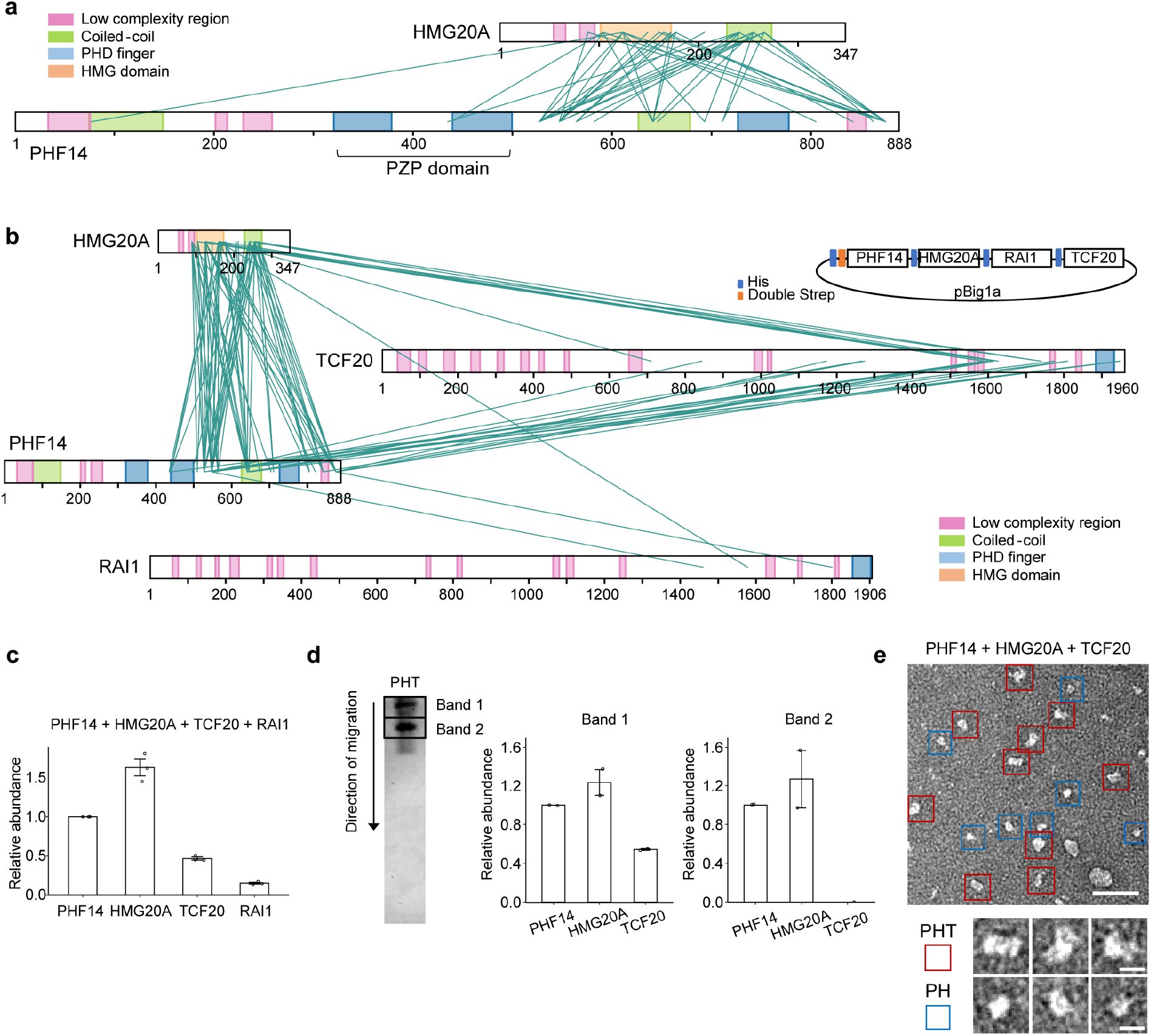
Interaction and stoichiometric analysis of the PHF14 complex purified in vitro. **a**, Visualization of XL-MS results using a cross-link viewer (xiNET) of PHF14 and HMG20A (PH). The green lines are indication of interactions. **b**, Visualization of XL-MS results using a cross-link viewer (xiNET) of PHF14, HMG20A, TCF20 and RAI1 and a schematic diagram of multigene expression vector used for expressing the proteins used in XL-MS. **c**, Relative abundances of the PHF14 complex proteins from iBAQ. Protein lysates are related to (**b**), n = 3. **d**, Non-denaturing electrophoresis gel of PHF14, HMG20A and TCF20 (PHT) and iBAQ results from the indicated native gel bands (n=2). **e**, Negative-stain EM micrographs of PH and PHT. Scale bar = 50 nm. Blue boxes and red boxes indicate PH and PHT, respectively. Three representative images of each proteins complex are shown below. Scale bar = 10 nm.

Next, we expressed all four proteins from a multigene DNA construct (Fig. 4b) in insect cells, and purified the PHF14 complex using ion exchange and size exclusion chromatography. The eluted fractions contained proteins with expected sizes corresponding to PHF14, HMG20A and RAI1/TCF20 (similar in size, over 200 kDa) (Extended Data Fig. 4c). Cross-linking MS showed that the interactions between PHF14 and HMG20A were conserved and the same regions interacted with the C-terminal region of TCF20 and to a lesser extent RAI1 (Fig. 4a, b, Extended Data Fig. 4d). The three statistically significant cross-links between PHF14-HMG20A and RAI1 were found towards its C-terminus (Fig. 4b), consistent with our domain analysis (Extended Data Fig. 2b). In addition, the purified complex had a lower abundance of RAI1 compared to TCF20 (Fig. 4c). The observation of intramolecular self-interaction cross-links between identical residues from PHF14 and HMG20A supports dimer formation of these two proteins, whereas such clear self-interaction cross-links were not observed for TCF20 or RAI1 (Extended Data Fig. 4e). Collectively, these data suggest that the binary core complex might form a ternary PHF14 complex with either TCF20 or RAI1.

Given the elevated levels of TCF20 relative to RAI1 in both ESCs (Fig. 1b) and the purified PHF14 complex, we focused on the ternary PHF14-HMG20A-TCF20 (PHT) complex. We co-purified the three proteins and confirmed their interactions (Extended Data Fig. 4f, g). Native gel electrophoresis showed two separate complexes (Fig. 4d). By MS intensity-based absolute quantification (iBAQ), a label-free protein quantification method, we identified the upper band as a PHF14-HMG20A-TCF20 complex with a 2:2:1 ratio and the lower band consisted of PHF14 and HMG20A only supporting a PHF14-HMG20A dimer as part of the complex.

As prior studies reported structural instability of purified macromolecular complexes^43^, we further determined homogeneity and dispersity of the purified PHT complex using negative stain electron microscopy (EM) (Fig. 4e). Despite a degree of heterogeneity, negative stain EM data displayed clear, discrete particles and a low level of aggregation at the concentration analyzed, indicating that the PHT complex readily forms and is maintained under our purification conditions.

### The PHF14 complex binds to double-stranded and four-way junction DNA

Given that PHF14 and HMG20A are rapidly recruited to DNA damage sites, we hypothesized that these proteins directly recognize DNA damage sites. We focused on HMG20A because it contains a high mobility group (HMG) box domain, a DNA-binding motif that has been reported to bind non-B-type DNA without sequence-specificity^44,45^. We performed electrophoretic mobility shift assays (EMSA) with single-stranded (ssDNA), double-stranded (dsDNA), a partially single-stranded four-way junction (ss4WJ) and a four-way junction (4WJ) synthetic DNA (Fig. 5a)^45^. The ss4WJ and 4WJ DNA (also called a Holliday junction) mimic non-B-type DNA structures which form during replication fork reversal and DNA repair. Using equimolar concentrations of DNA, we found that HMG20A bound strongly to dsDNA, ss4WJ and 4WJ DNAs with weakest binding to ssDNA (Fig. 5b and Extended Data Fig. 5a).

**Fig. 5:**
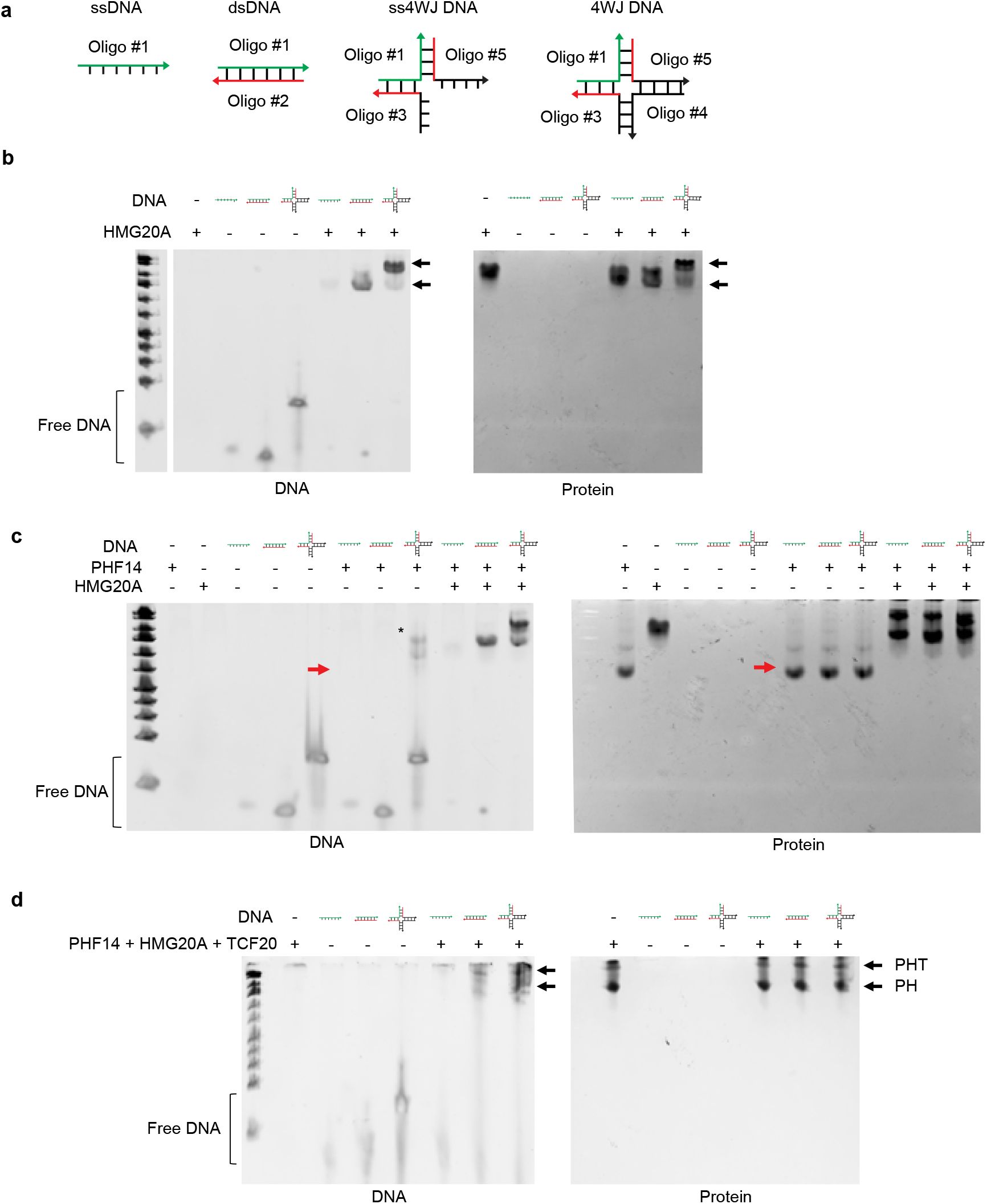
PHF14 complex binds to dsDNA and four-way junction DNA mediated by HMG20A, but PHF14 does not bind to DNA alone. **a**, Visualization of DNA shapes used in EMSA. The nucleotide sequences were adopted from Bianchi et al., 1989. The identical sequences among the different DNAs are in the same color (green and red). The arrows indicate the direction of DNAs from 5’ to 3’. **b**, EMSA with HMG20A proteins on a non-denaturing polyacrylamide gel. HMG20A proteins were incubated with different shapes of DNA indicated above of each panel and stained for both DNA and proteins. Arrows indicate positions of DNA bands bound by HMG20A protein showing shift in mobility. **c**, EMSA with PHF14 and HMG20A proteins, incubated together with different shapes of DNA, on a non-denaturing polyacrylamide gel. Arrows indicate the position of PHF14 protein. Asterisk indicates non-specific bands. **d**, Purified proteins from Fig. 4e incubated with DNA of different conformations. Arrows indicate the locations of the band with just PHF14 and HMG20A (PH) and all three proteins (PHF14, HMG20A and TCF20; PHT). In all experiments, DNA concentration was 1 μM and protein concentrations were 1 μM of each protein for (**b**) and (**c**). Oligonucleotide sequences are listed in Supplementary Table 2.

To further characterize the DNA-binding properties of HMG20A, we performed HMG20A titrations with a fixed amount of DNA (Extended Data Fig. 5a and Fig. 5b). All of the 4WJ DNA was bound even with the lowest concentration of HMG20A; in contrast, free ss4WJ DNA was still visible at the lowest HMG20A concentration and free dsDNA was observed even at the highest HMG20A concentration (Extended Data Fig. 5a). These data suggest that HMG20A has different affinity for DNA depending on its structure with stronger affinity to 4WJ DNA.

Next, we determined whether PHF14 was capable of binding DNA either alone or with HMG20A. PHF14 alone did not bind these DNA structures, as free DNAs were visible and the shifted 4WJ DNA signal was non-specific and not matched to the PHF14 protein on the gel (red arrow, Fig. 5c). HMG20A incubated with increasing concentrations of PHF14 in the absence of DNA resulted in two subpopulations with no free PHF14 remaining (Extended Data Fig. 5b), consistent with a strong protein-protein interaction. By EMSA, we found that the PHF14-HMG20A complex bound dsDNA and 4WJ, but minimally bound ssDNA (Fig. 5c). We performed EMSA with the purified ternary PHT complex and our data revealed that both the PH (lower band) and PHT (upper band) populations (as identified in Fig. 4e) bind dsDNA and 4WJ, but not ssDNA, suggesting that the ternary complex can indeed form on DNA (Fig. 5d).

### The PHF14 complex forms DNA-containing phase separated condensates

Thus far, our experiments revealed that HMG20A facilitates the dsDNA and 4WJ binding of the PHF14 complex, consistent with a possible role in the DNA damage-sensing first step of recruitment, but we still lacked an understanding of the function of the complex at DNA damage sites, particularly in terms of TCF20 and RAI1 recruitment to the PHF14 complex. TCF20 and RAI1 have annotated domains only in their C-termini, despite their large size (1960 and 1906 amino acids, respectively). To better understand the biophysical properties of the rest of these proteins, we used several domain search tools^46^ and found that all four proteins contain several intrinsically disordered regions (IDR) with low hydrophobicity (Fig. 6a and Extended Data Fig. 6a). TCF20 and RAI1 also include predicted prion-like amino acid composition (PLAAC)^47^ near their N-termini and contain several regions with predicted planar pi-pi contacts^48^. Such IDRs have been identified in phase separating proteins, such as FUS^49,50^. Recent studies have reported that some DDR proteins form biomolecular condensates and that the formation of DNA damage foci involves liquid-liquid phase separation^51,52^. Thus, we hypothesized that the PHF14 complex might function at DNA damage sites by forming biomolecular condensates (Fig. 6b).

**Fig. 6:**
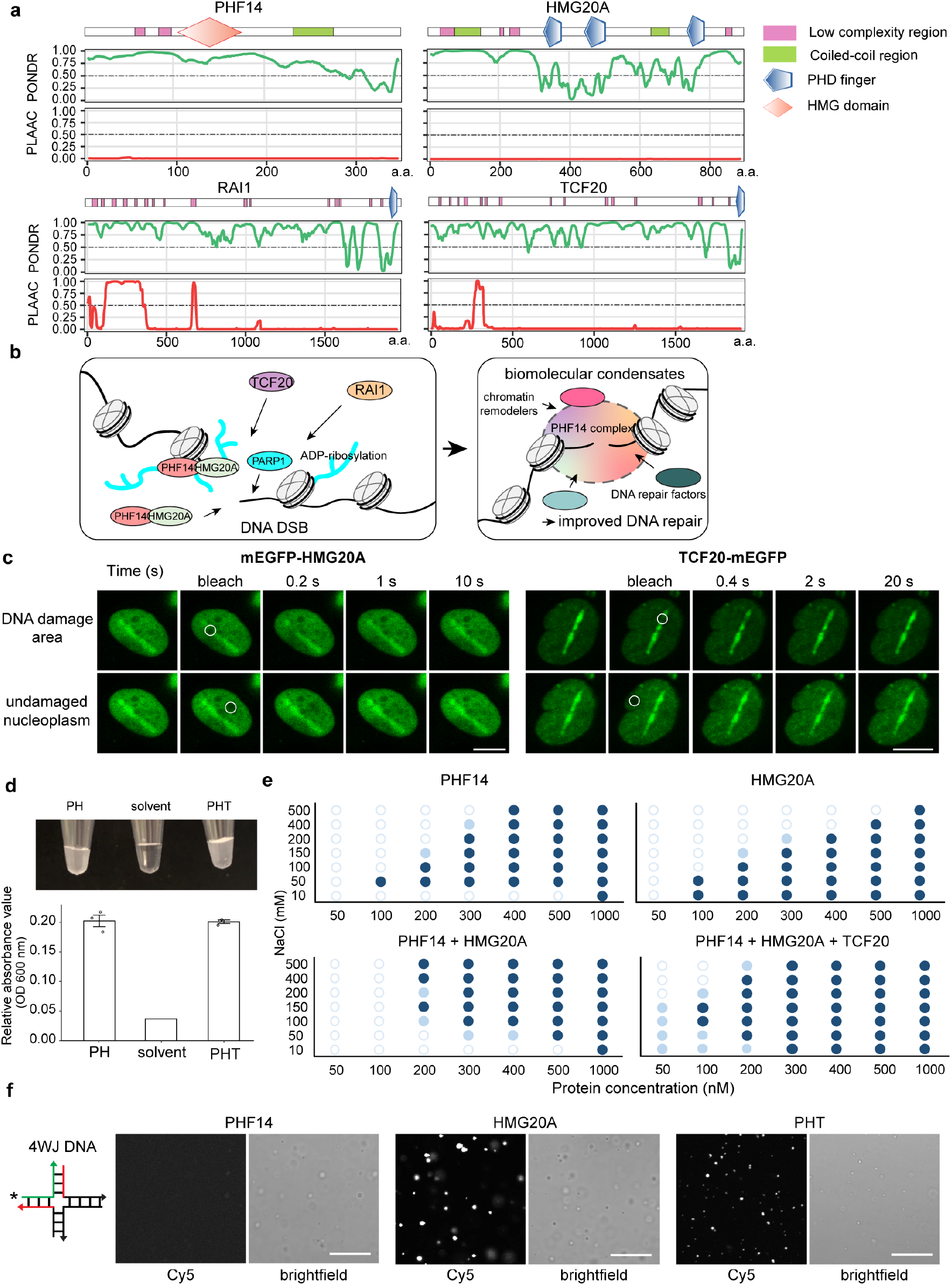
PHF14 complex undergoes phase separation in vitro and the formation of droplets is enhanced by TCF20. **a**, Predictions of IDRs from amino acid sequences of PHF14 complex. Each upper-most panel indicates both predicted protein domains and IDRs from Simple Modular Architecture Research Tool (SMART), then Predictor of Natural Disordered Regions (PONDR) and Prion-Like Amino Acid Composition (PLAAC). Scores greater than 0.5 are indicative of disorder propensity. Values on the x-axis correspond to amino acid residue number of the corresponding protein. **b**, Graphical summary of the phase separation hypothesis. PHF14 and HMG20A form a binary complex that is recruited to DNA damage sites as a first step at the same speed as PARP1. PARP1 PARylates (shown in turquoise) both of them and its other targets at DNA damage sites. TCF20 and RAI1 are recruited later, PARylation-dependently and a tertiary or quaternary protein complex is assembled at the damage sites. The local increase in protein concentration triggers the formation of biomolecular condensates that separate the damage sites from the surrounding nucleoplasm and facilitate other repair factor recruitment. **c**, Confocal microscopy images of FRAP in U2OS cells transfected with mEGFP-HMG20A or TCF20-mEGFP. FRAP areas were bleached with 488 nm laser at the indicated locations. DNA damage was induced with 355 nm laser and then proteins were allowed to reach maximal accumulation before FRAP, where applicable. Scale bars = 10 μm. **d,** Comparison of turbidity changes of the PHF14 complex after phase separation (PH – PHF14 and HMG20A, PHT – PHF14, HMG20A and TCF20). Quantification was done by measuring OD 600 nm. **e**, Phase diagrams of proteins indicated in each diagram, with changes in protein concentration on the x-axis and NaCl concentration on the y-axis. The experiment was repeated three times and representative results are shown. Dark blue = clear droplets, light blue = some droplets, white = no droplets. **f**, Confocal images of individual proteins and the three-protein complex (PHT) after undergoing phase separation with the addition of Cy5-labeled 4WJ DNA. Scale bars = 20 μm.

We examined the dynamics of the PHF14 complex accumulated areas at DNA damage sites in live cells using fluorescence recovery after photobleaching (FRAP) analysis. mEGFP-HMG20A and TCF20-mEGFP showed a rapid recovery at damaged DNA, with a t_50_ of 0.8 s for HMG20A and 2.3 s for TCF20, similar to the FRAP speed of undamaged nucleoplasmic regions (Fig. 6c, Extended Fig. 6b). These fast recovery speeds indicate that the PHF14 complex is a dynamic component of the DNA damage response, capable of macromolecule exchange, unlike solid-like aggregates. These results are consistent with the properties of biomolecular condensates within cells.

To determine whether the PHF14 complex can form phase separated condensates in vitro, we examined the purified binary (PHF14-HMG20A) and ternary (PHF14-HMG20A-TCF20) complexes. Under a widefield microscope, we observed spherical droplets with dynamic movement right after mixing, behaving like liquid when they touch a glass surface in physiological salt conditions (150 mM NaCl) and with 5% PEG as a crowding reagent (Extended Data Fig. 6c, d). Moreover, turbidity measurement of the proteins in solution showed that the protein solutions with these droplets had much higher optical density compared to the solvent (Fig. 6d), consistent with biomolecular condensates. We also found that either PHF14 or HMG20A alone formed droplets, with each component and the PHF14-HMG20A complex displaying a distinct optimal salt concentration (Fig. 6e). PHF14-HMG20A droplets were more resistant to high salt concentrations than single proteins alone, consistent with extensive multivalent interactions. Notably, the presence of TCF20 enhanced phase separation at lower concentrations of PHF14-HMG20A (Fig. 6e).

To investigate whether the PHF14 complex can still form condensates when bound to DNA, we added Cy5-labeled 4WJ DNA to the ternary PHF14 complex (PHF14-HMG20A-TCF20) and performed confocal microscopy. We observed the formation of droplets that all contained Cy5-DNA while still maintaining liquid-like properties when they touch a glass surface (Extended Data Fig. 6d). Our EMSA analysis suggests that PHF14 binds to DNA through HMG20A (Fig. 5), so we examined whether HMG20A was necessary for the incorporation of Cy5-labeled 4WJ DNA into the droplets. Indeed, HMG20A droplets but not PHF14 droplets contained DNA (Fig. 6f), suggesting that HMG20A mediates the inclusion of 4WJ DNA into the phase-separated PHF14 complex.

Although the presence of nucleic acids can alter the formation of biomolecular condensates ^53,54^, we found that neither the presence nor shape of DNA affected the efficiency of condensate formation for the PHF14-HMG20A complex at physiological salt concentration (Extended Data Fig. 6d). Therefore, condensate formation is likely driven by the local increase in protein concentration that in the cells is achieved by accumulation at DNA damage sites. Altogether, we propose that the PHF14 complex recruited to DNA damage areas forms dynamic biological condensates with DNA.

### TCF20 affects the properties of the PHF14 complex biomolecular condensates

Because TCF20 enhanced condensate formation at lower protein concentrations (Fig. 6e), we examined the biophysical properties of the condensates and whether these are affected by the presence of TCF20. We found that the Cy5-4WJ DNA containing droplets were dynamic in their movement and fused with each other (Fig. 7a). Using FRAP, we observed active exchange of DNA with the solution outside of the droplets (PHT: t_50_ 6 s, HMG20A: t_50_ 66 s) and also within the droplets (PHT: t_50_ 24 s, HMG20A: 36 s) (Fig. 7b and Extended Data Fig. 7a). Notably, the FRAP time-series showed faster recovery with PHT proteins (t_50_ of 6 s) compared to HMG20A alone (t_50_ of 66 s), raising the possibility that TCF20 regulates the dynamics of condensates.

**Fig. 7:**
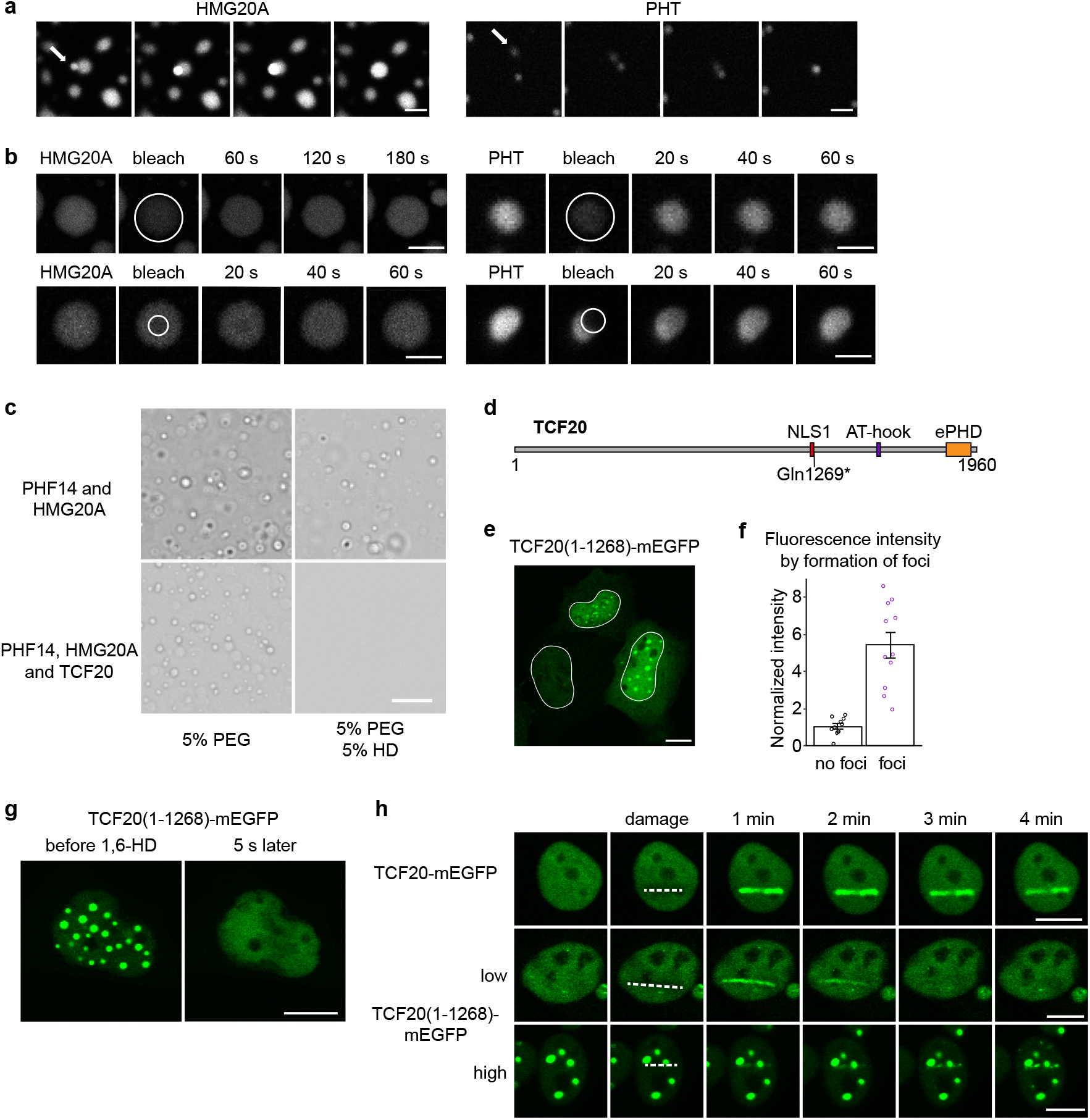
TCF20 makes the PHF14 complex condensates more dynamic. A patient mutation-truncated TCF20 can still form condensates and be recruited to damage sites, but is not maintained there. **a**, Confocal microscopy images of Cy5-labeled dsDNA in phase separated protein droplets showing droplets merging (indicated by the white arrow). Scale bar = 10 μm. **b**, Whole-droplet and partial FRAP of the phase separated droplets for Cy5-labeled DNA. Scale bars = 5 μm. **c**, Phase contrast images of condensates that are formed with indicated proteins with and without 1,6-hexanediol. 300 nM of proteins were used. Images were taken with a widefield Nikon Ti-E microscope. Scale bar = 20 μm. **d**, Schematic of TCF20 protein. NLS1 (aa 1254-1268) precedes the Gln1269* patient mutation, explaining the nuclear localization of the TCF20(1-1268) construct. **e**, TCF20(1-1268)-mEGFP transfected into U2OS cells. White outlines indicate nuclear borders. Scale bar = 10 μm. **f**, Correlation between fluorescent signal intensity in TCF20(1-1268)-mEGFP transfections with the formation of foci. Data from 3 frames, normalized within frame with the average signal from cells without foci set to 1. **g**, TCF20(1-1268)-mEGFP before and immediately after the addition of 1,6-hexanediol and digitonin to final concentrations of 5% and 5 μg/mL, respectively. Imaged with 5 s frame interval. Scale bar = 10 μm. **h**, Confocal microscope images of full length TCF20-mEGFP and TCF20(1-1268)-mEGFP accumulation at DNA damage sites after damage induction with 355 nm laser. Scale bars = 10 μm.

This was further supported by the effects of 1,6-hexanediol (HD) on the droplets in vitro. We found that HD completely abolished the condensates formed by the ternary complex while the binary complex consisting of HMG20A and PHF14 was only partly affected (Fig. 7c). As HD dissolves liquid-like assemblies but not solid-like states^50^, these results suggest that the recruitment of TCF20 to PHF14-HMG20A complexes at DNA damage sites facilitates the formation of the liquid-like phase-separated compartments.

### Truncated TCF20(1-1268) forms aggregates and is not maintained at DNA damage sites

Given that TCF20 changes the biophysical properties of the PHF14 complex droplets, we hypothesized that TCF20 helps to establish and maintain phase-separated DNA damage regions and mutations in TCF20 might affect this process. To determine whether TCF20 mutations reported in patients with neurodevelopmental disorders alter TCF20 localization to damaged DNA, we generated a truncated TCF20(1-1268)-mEGFP that mimics a nonsense mutation reported in a patient^5^. This mutant TCF20 lacks the C-terminus, where the known structured domains and most of the interactions with PHF14-HMG20A are located (Fig. 4b, 7d and Extended Data Fig. 4g), but the first nuclear localization signal (NLS) and the predicted IDRs are still present, including the N-terminal region with prion-like amino acid composition (Fig. 6a and Extended Data Fig. 6a).

In contrast with the full length TCF20-mEGFP (Extended Data Fig. 7b), TCF20(1-1268)-mEGFP aggregated into large foci in U2OS cells (Fig. 7e and Extended Data Fig. 7c). Nuclei with foci had a higher level of fluorescence, whereas nuclei with no foci and diffuse nucleoplasmic signal had lower fluorescence intensity (Fig. 7f), suggesting that foci formation is concentration-dependent. The large foci formed by the truncated TCF20 regardless of DNA damage generation were dissociated upon addition of 1,6-hexanediol (Fig. 7g), implying that these droplets exist in a liquid-like state.

After inducing DNA damage, the low expression cells without foci showed recruitment of TCF20(1-1268)-mEGFP to DNA damage sites. However, it was not maintained and instead was lost in 3 minutes (Fig. 7h and Extended Data Fig. 7f). TCF20(1-1268)-mEGFP recruitment to DNA damage sites was also observed in the cells with foci, but it quickly condensed into individual foci along the damage line (Fig. 7h and Extended Data Fig. 7g). Thus, the C-terminus of TCF20 that is often lost in patients with TCF20 developmental syndrome is not necessary for its initial recruitment to damage sites or its ability to form phase separated condensates, but, likely through its interactions with the PHF14-HMG20A core complex (Fig. 4), it is required to maintain its correct localization and for the regulation of the condensates.

## Discussion

Here, we demonstrate that the PHF14 complex containing HMG20A and intellectual disability-related paralogs RAI1 or TCF20 is a novel regulator of the DDR. Our results support a stepwise recruitment mechanism for the PHF14 complex at DNA damage sites with the HMG20A-PHF14 dimer binding first. TCF20 and RAI1 are subsequently recruited and might be mutually exclusive subunits. We also observed that PHF14-HMG20A-TCF20 form phase separated condensates in vitro. The condensates are spherical, fuse with each other and show fast fluorescence recovery after photobleaching. HMG20A and PHF14 can phase separate individually or together, and TCF20 allows these proteins to form condensates at lower concentrations. Most interactions between PHF14-HMG20A and TCF20 map to the C-terminal part of TCF20. A TCF20 construct corresponding to a patient mutation missing the C-terminus appears to form droplets in cells at concentrations where full length TCF20 shows diffuse nucleoplasmic signal. The mutant protein is still recruited to DNA damage sites, but quickly coalesces into droplets. Thus, the C-terminus of TCF20 (and RAI1) interacts with PHF14 and HMG20A to form a protein complex and may play a role in regulating condensate formation.

Several phase-separating proteins, such as FUS, have been reported to be involved in DNA repair, possibly through the recruitment of repair factors into condensed hubs with broken DNA. In addition, DDR factors have been recently shown to also have phase separating properties themselves, such as 53BP1 and Rad52^52,55^, or to regulate the formation of droplets, for example, PARylation has been reported to nucleate intracellular liquid demixing^51,56^. Interestingly, TCF20 and RAI1 recruitment to DNA damage sites is also dependent on PARylation. It is possible that the low-complexity regions of these proteins can interact with PAR, as reported for other intrinsically disordered proteins^51,56^. We envision that the PHF14 complex collaborates with other phase-separating proteins to ensure the efficacy of DNA repair.

Chromatin remodeler complexes are known to be recruited to DNA damage sites as an early step^57,58^. Notably, the PHF14 complex is recruited even faster than remodeler proteins, and HMG20A and PHF14 localize to DNA damage sites at a comparable speed to PARP1, which is one of the earliest responders. PARP1 directly senses DNA damage^59^ and based on the speed alone, we speculate that the PHF14 complex might do the same, possibly through the DNA-binding ability of Hmg20a. Members of the NuRD complex, HDAC1 and HDAC2 in particular, are recruited to DNA damage sites with a t_50_ of 3.5-5 s and are maintained there for over an hour^60^. We found that NuRD complex components interact with Phf14 in response to DNA damage, suggesting that the PHF14 complex might help recruit or maintain the localization of the chromatin remodelers at damage sites.

We show that loss of Phf14 leads to defects in cell cycle progression and cell proliferation, particularly in NPCs. Disrupted cell cycle and proliferation in NPCs are the underlying cause of several developmental delay syndromes^61^. At the early stage of neurodevelopment, NPCs undergo massive expansion and during this time precise control of genome replication is essential. When replication errors and stress occur that induce DSBs, DNA damage response and DNA repair must operate accurately on target sites. Temporal control of cell migration is also critical for brain development, so even subtle delays in cell cycle progression and cell division caused by DNA damage that is eventually resolved could strongly impact neurodevelopment. Thus, the proliferation defects in *Phf14* KO NPCs might explain the neurological phenotypes observed in patients with mutations in the PHF14 complex members. These cell cycle defects could be in part caused by decreased DNA repair efficiency in *Phf14* KO cells, which also have increased susceptibility to DNA damaging agents. The PHF14-HMG20A binding to 4WJ and ss4WJ DNA might contribute directly to the protection of stalled replication forks that we found to be defective in *Phf14* KO cells, as these structures resemble reversed and stalled replication forks, respectively, and are also formed as an intermediate step in DNA repair^55,62^. Further structural studies of the PHF14 complex with DNA might shed light on the molecular mechanism.

Mutations that disrupt DDR pathways cause diseases with neurological symptoms and are usually accompanied by increased cancer risk or immunological deficiencies, e.g., Nijmegen breakage syndrome associated with mutations in NBS1, part of the MRN complex that processes DSBs in both HR and NHEJ pathways^63^, and mutations in the NHEJ ligase LIG4^64^. Many patients with these syndromes exhibit microcephaly, which indicates that the impaired DNA repair strongly affects NPC proliferation. Recent reports have also identified cases that mainly affect the nervous system, as exemplified by core histone mutation (e.g. H4K91R) that causes intellectual disability through genome instability^65^. Thus, it is possible that less severe defects in DDR primarily affect the highly sensitive nervous system and the impaired function of the PHF14 complex in DDR could be the cause of the symptoms experienced by RAI1 and TCF20 mutation patients.

## Competing interests

Authors declare no competing interests.

## Data and materials availability

Next-Generation Sequencing datasets (mRNA-seq) have been deposited in the ArrayExpress database and the mass spectrometry proteomic data have been deposited to the ProteomeXchange Consortium via the PRIDE^66^ partner repository. The data will be made publicly available upon peer-reviewed publication.

## Acknowledgments

We thank the staff at the European Molecular Biology Laboratory (EMBL) Advanced Light Microscopy Facility, the Genomics Core Facility, the Flow Cytometry Core Facility, the Protein Expression and Purification Core Facility, and the Proteomics Core Facility for training, consultation, sample preparation and data generation. We thank C. Girardot and the Genome Biology Computational Support as well as the EMBL Centre for Bioimage Analysis for their assistance in data analysis, and the Noh laboratory for reading the manuscript and for helpful discussions. This work is supported by the EMBL predoctoral fund (to K.M.N.), the DFG fund (SPP 1738 to K.M.N.), the EIPOD postdoctoral fund under Marie Sklodowska-Curie Action (664726, to I.-Y.H. and H.K.H.F.).

---

**Extended Data Fig. 1:**
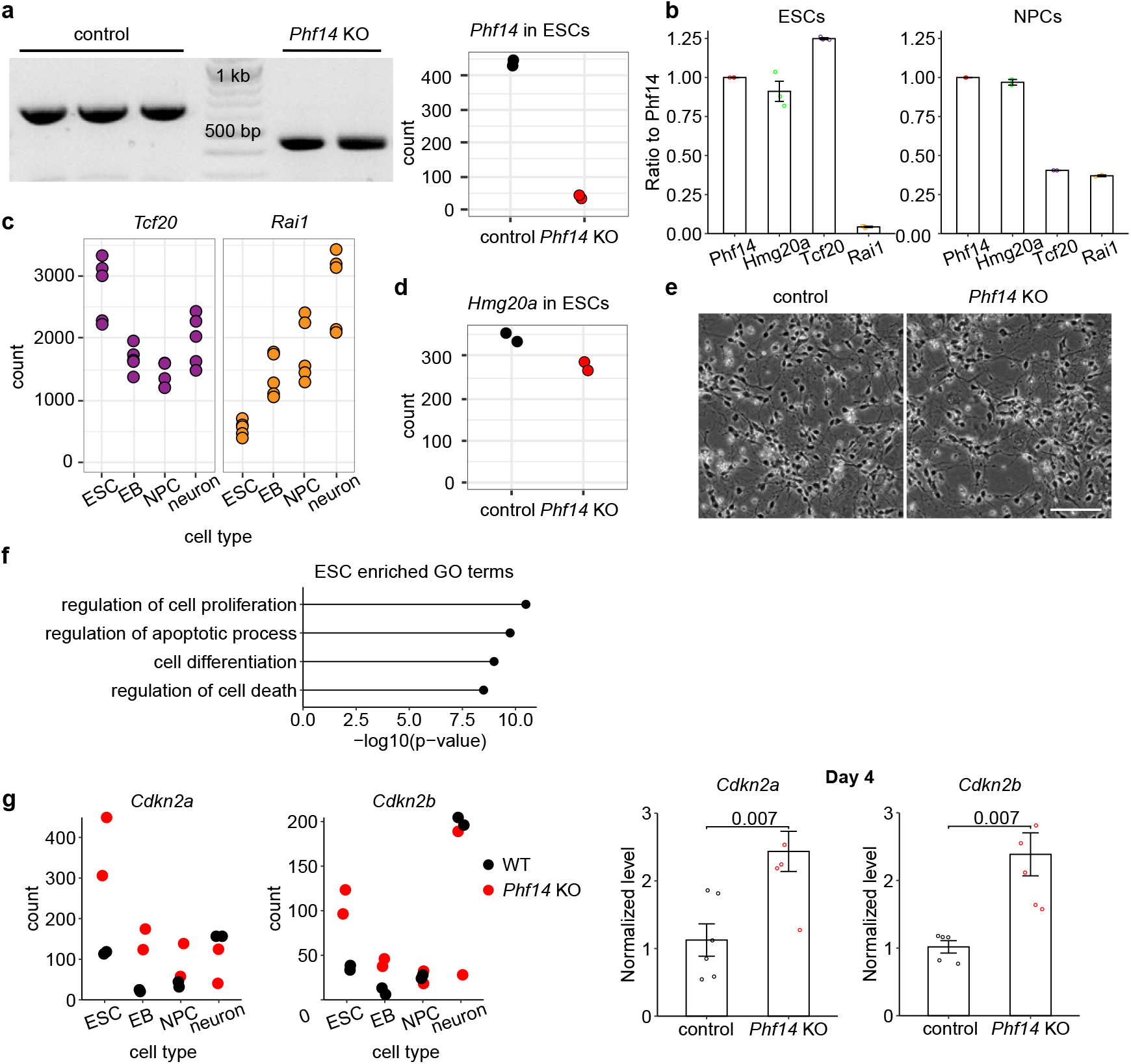
Phf14 forms a complex with Hmg20a, Tcf20 and Rai1; *Phf14* KO cells exhibit cell proliferation defects. **a,** Left: PCR amplicons on agarose gel around exon 5 of Phf14 (wild-type: 654 bp, KO: around 460 bp). Right: *Phf14* mRNA-levels (library size normalized counts mapping to the gene) in *Phf14* KO and WT ESCs. **b**, Approximate stoichiometry of the Phf14 complex from pulldowns with Phf14 antibody from ESCs and NPCs. Calculated as 10^(top3-value)*signal_sum(given TMT lane)/signal_sum(all TMT lines). n = 3 (ES cells) and n = 2 (NPCs) **c**, mRNA levels (library size normalized counts mapping to the gene) of Tcf20 and Rai1 in control cells during differentiation (D0 – ESCs, D4 – embryoid bodies, D8 – NPCs, D12 – neurons). **d,** Hmg20a mRNA-levels (library size normalized counts mapping to the gene) in *Phf14* KO and WT ESCs. **e**, Widefield microscope images of control and *Phf14* KO neurons. Scale bar = 100 μm. **f**, GO enrichment analysis for genes differentially expressed in *Phf14* KOs compared to controls in ESCs. **g** Cdkn2 and Cdkn2b levels from mRNA-sequencing (library size normalized counts mapping to the gene) and RT-qPCR at day 4 (embryoid body stage).

**Extended Data Fig. 2:**
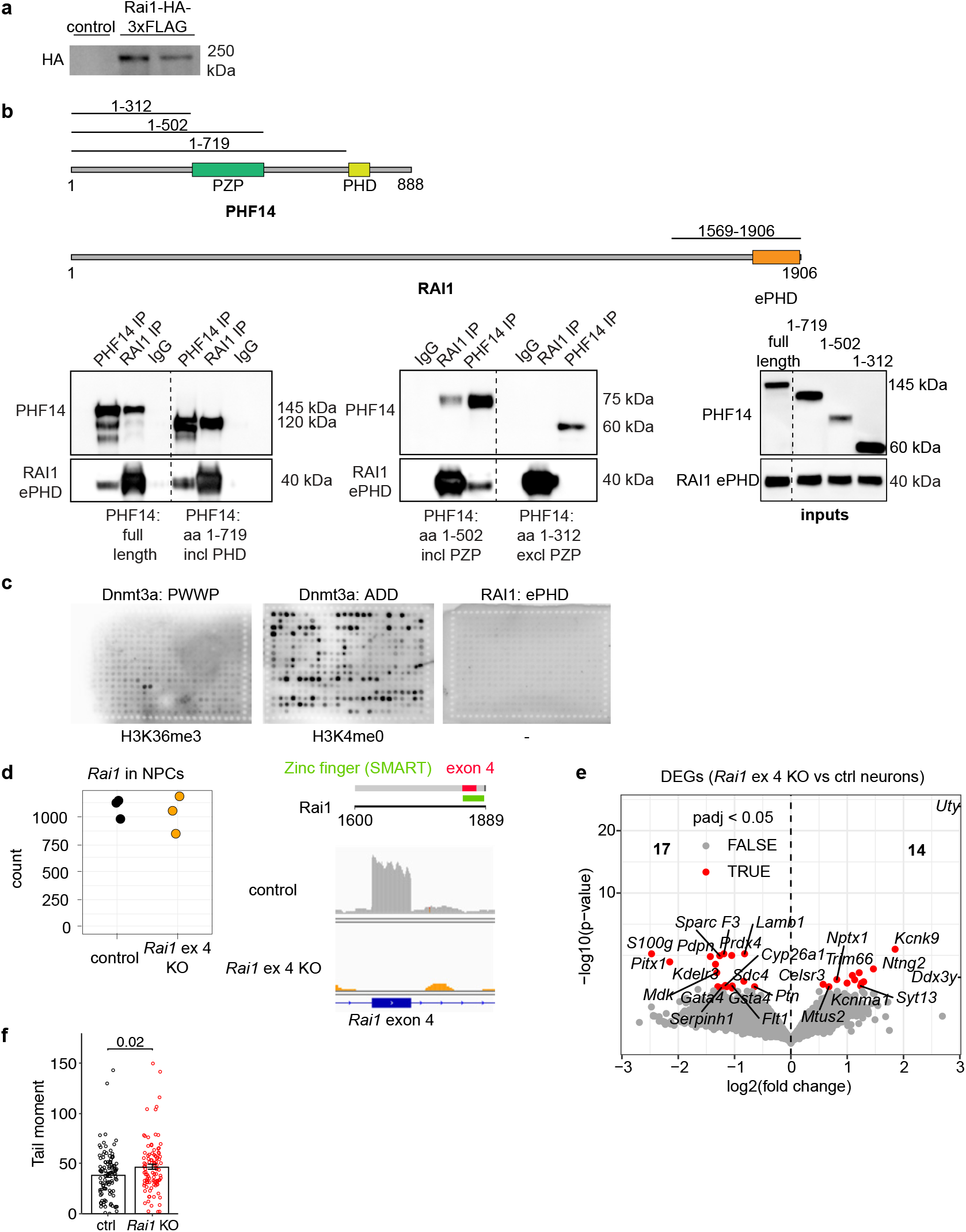
The ePHD of RAI1 does not bind histone tails but instead interacts with PHF14. **a,** Western blot showing HA-tagged Rai1 at the expected size (slightly below 250 kDa) in single-cell selected clonal lines. **b**, Co-immunoprecipitation and inputs of RAI1 ePHD and PHF14 constructs detected by Western blot in HEK293T cells. Constructs: HA-FLAG-RAI1(1569-1906), myc-PHF14(1-888), myc-PHF14(1-719), myc-PHF14(1-502) and myc-PHF14(1-312). **c**, MODified histone peptide array of purified ePHD domain of Rai1. Dnmt3a PWWP and ADD domains were used as a positive control. Representative images from 2 replicates. **d**, *Rai1* mRNA normalized counts from mRNA-seq data and IGV browser views of the reads mapped to *Rai1* gene with the deleted exon 4 highlighted. **e**, Volcano plot from mRNA-seq analysis with DESeq2 showing all differentially expressed genes in neurons with adj. p-value < 0.05. (3 independent replicates in both groups) (Full list of identified DEGs provided in Source data.) **f**, Comet tail moments after etoposide treatment, quantified using CometScore 2.0 software.

**Extended Data Fig. 3:**
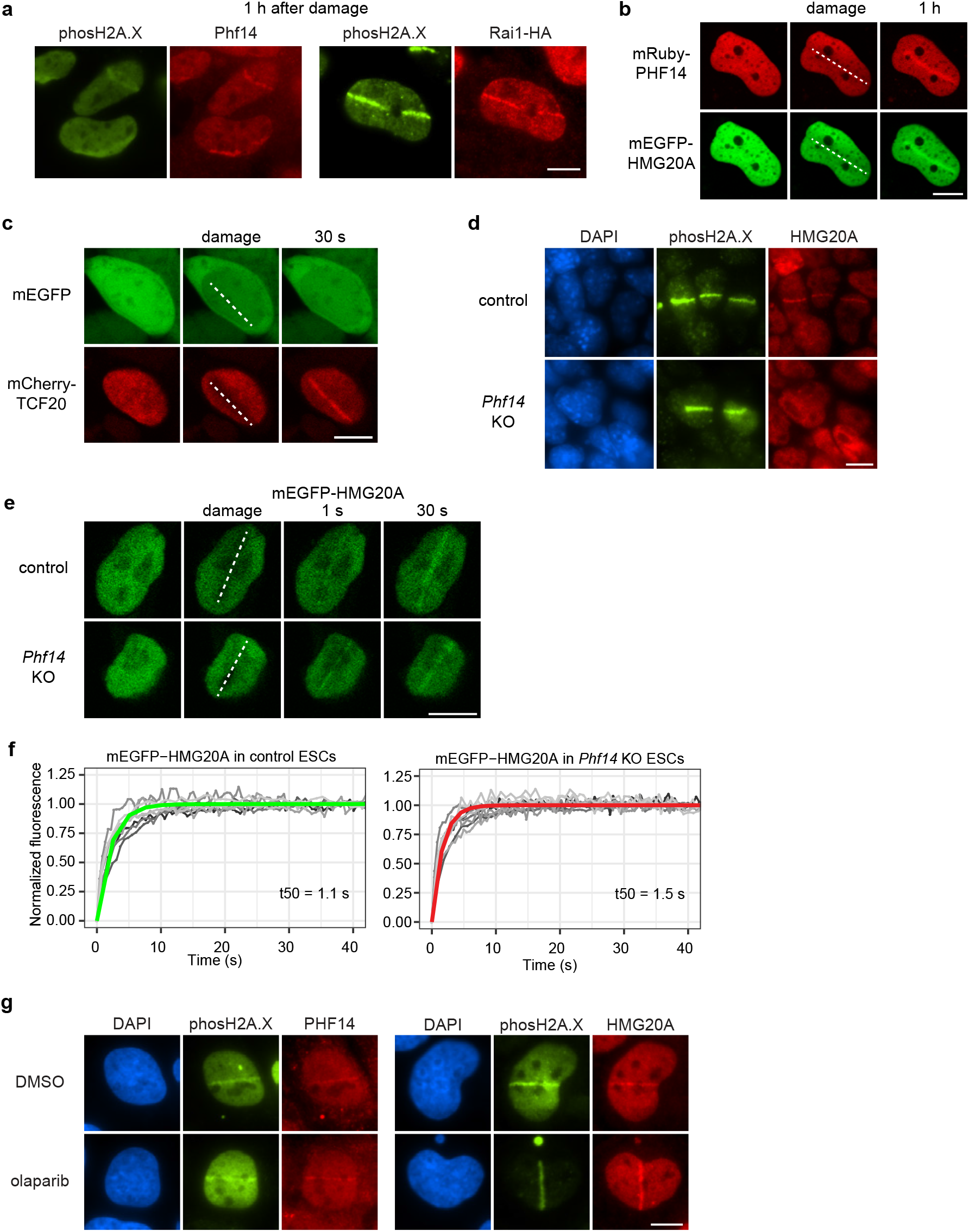
The PHF14 complex is recruited to DNA damage sites. **a**, Immunofluorescence staining in monolayer-differentiated NPCs after laser microirradiation. Anti-phospho-histone H2A.X (Ser139) signal serves as a DNA damage marker. Cells were fixed 1 h after DNA damage was generated. Laser microirradiation in transiently transfected U2OS cells, 1 hour after DNA damage induction. **b**, Live-cell imaging of mEGFP-HMG20A and mRuby-PHF14 transiently transfected into U2OS cells. 1 hour after damage induction. **c**, Live-cell imaging of mEGFP and TCF20-mCherry (transient co-transfection into U2OS cells). DNA damage was induced with 355 nm laser at the indicated location. **d**, Immunofluorescence staining in control and *Phf14* KO ESCs after laser microirradiation. **e**, Live-cell imaging of mEGFP-HMG20A, transiently transfected into control and *Phf14* KO ESCs. DNA damage was induced with 355 nm laser at the indicated location. **f**, Quantified recruitment kinetics for mEGFP-HMG20A in control and *Phf14* KO ESCs. **g**, Immunofluorescence staining in U2OS cells after laser microirradiation. DNA damage was induced after treatment with 1 μM olaparib for 1 h or equivalent amount of DMSO. All scale bars = 10 μm.

**Extended Data Fig. 4:**
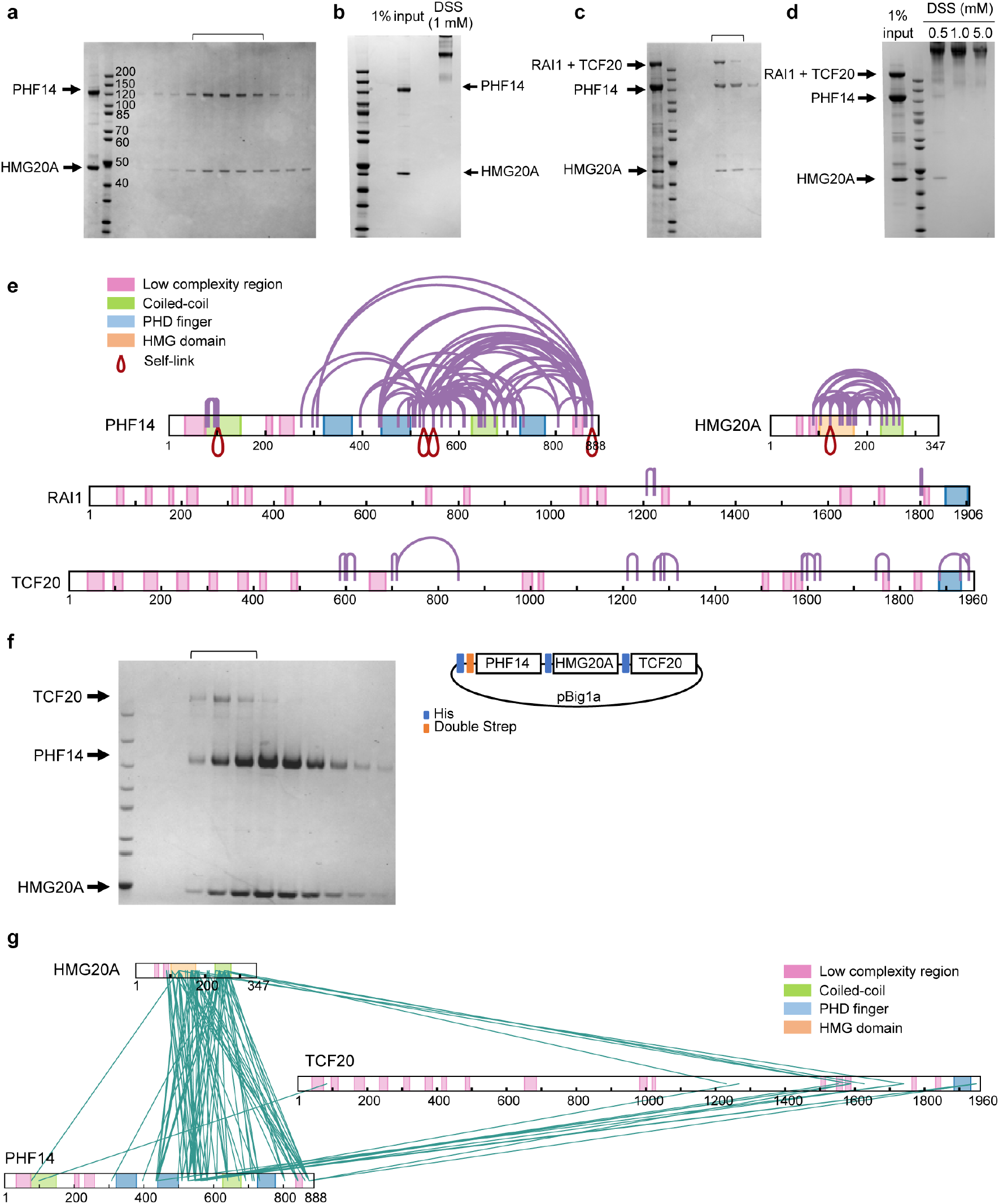
Details of Protein Purification and XL-MS of PHF14 complex. **a**, Protein gel staining of gel filtration of the purified proteins from co-lysis of PHF14 and HMG20A expressing insect cells. Collected fractions are indicated with the square bracket. **b**, SDS-PAGE input PHF14 and HMG20A, and after cross-linking with DSS. **c**, Protein gel staining (left) and graph ^67^ of gel filtration of the purified complex with PHF14, HMG20A, TCF20 and RAI1. Final eluted fractions were confirmed by SDS-PAGE for expression of the PHF14 complex. Collected fractions are indicated with the square bracket. **d**, Purified proteins from (**c**) were tested for optimal cross-linker concentration, DSS. SDS-PAGE of cross-linked and input complex. **e**, Intra-protein interactions of XL-MS result visualized by xiNET. Related to Fig. 4c. **f**, Schematic diagram of PHT multigene expression cassette and protein gel staining from the final purified proteins from gel filtration. **g**, XL-MS results showing the interactions of PHT complex.

**Extended Data Fig. 5:**
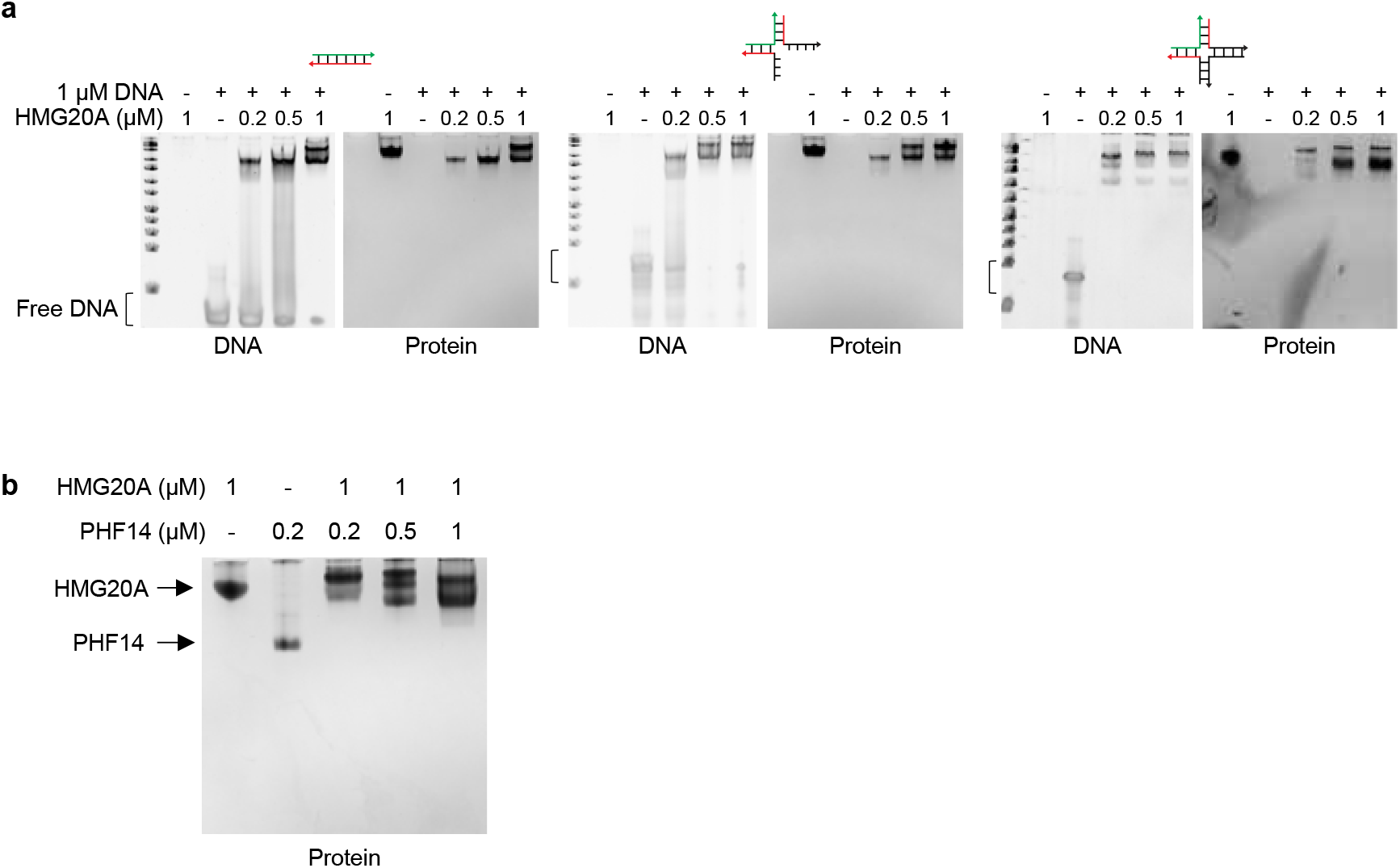
Non-denaturing gel electrophoresis results of HMG20A and PHF14. **a**, Purified HMG20A protein incubated with the DNA indicated each lane. Protein concentrations are as indicated. **b**, Individual proteins of PHF14 and HMG20A purified separately and analyzed by non-denaturing gel electrophoresis separately and mixed together.

**Extended Data Fig. 6:**
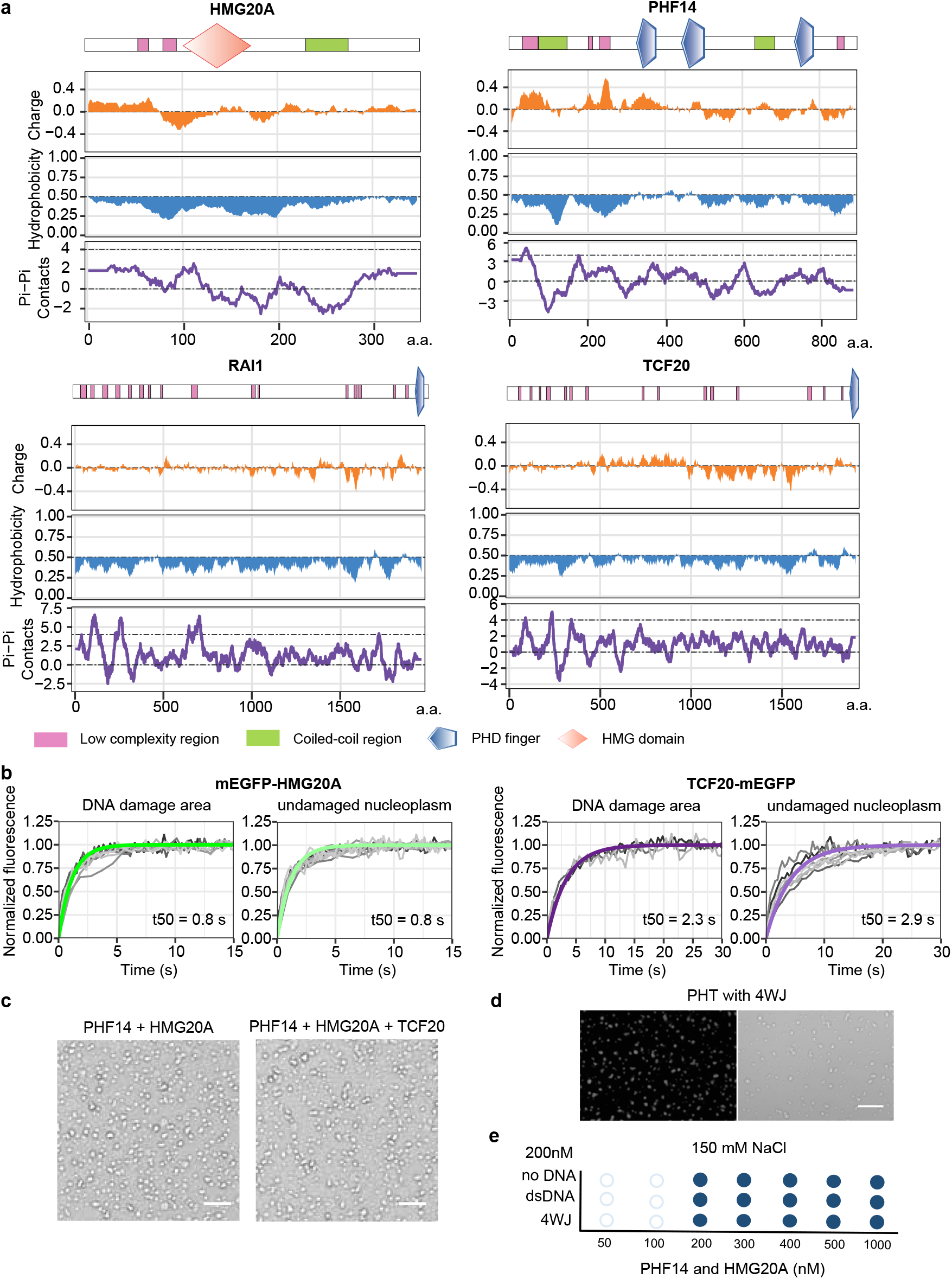
Biochemical properties of the PHF14 complex. **a**, Charge, hydrophobicity and pi-pi contact predictions. For pi-pi contacts, greater than 4 is indicative of phase separation. Values on the x-axis correspond to amino acid residue number of the corresponding protein. **b**, Quantified recovery kinetics from FRAP experiments in Fig. 6c. **c**, Phase contrast images of condensates that are formed with indicated proteins and 10% of PEG. Images were taken with a widefield Nikon Ti-E microscope **d**, Glass surface wetting assay of PHT proteins with Cy5-labeled DNA from confocal microscope. Scale bar = 20 μm. **e**, Phase diagram of PHF14 and HMG20A proteins with different shapes of DNA (200 nM) (y-axis) and increasing protein concentration (x-axis) at physiological salt concentration (150 mM NaCl). Dark blue = clear droplets, light blue = some droplets, white = no droplets.

**Extended Data Fig. 7:**
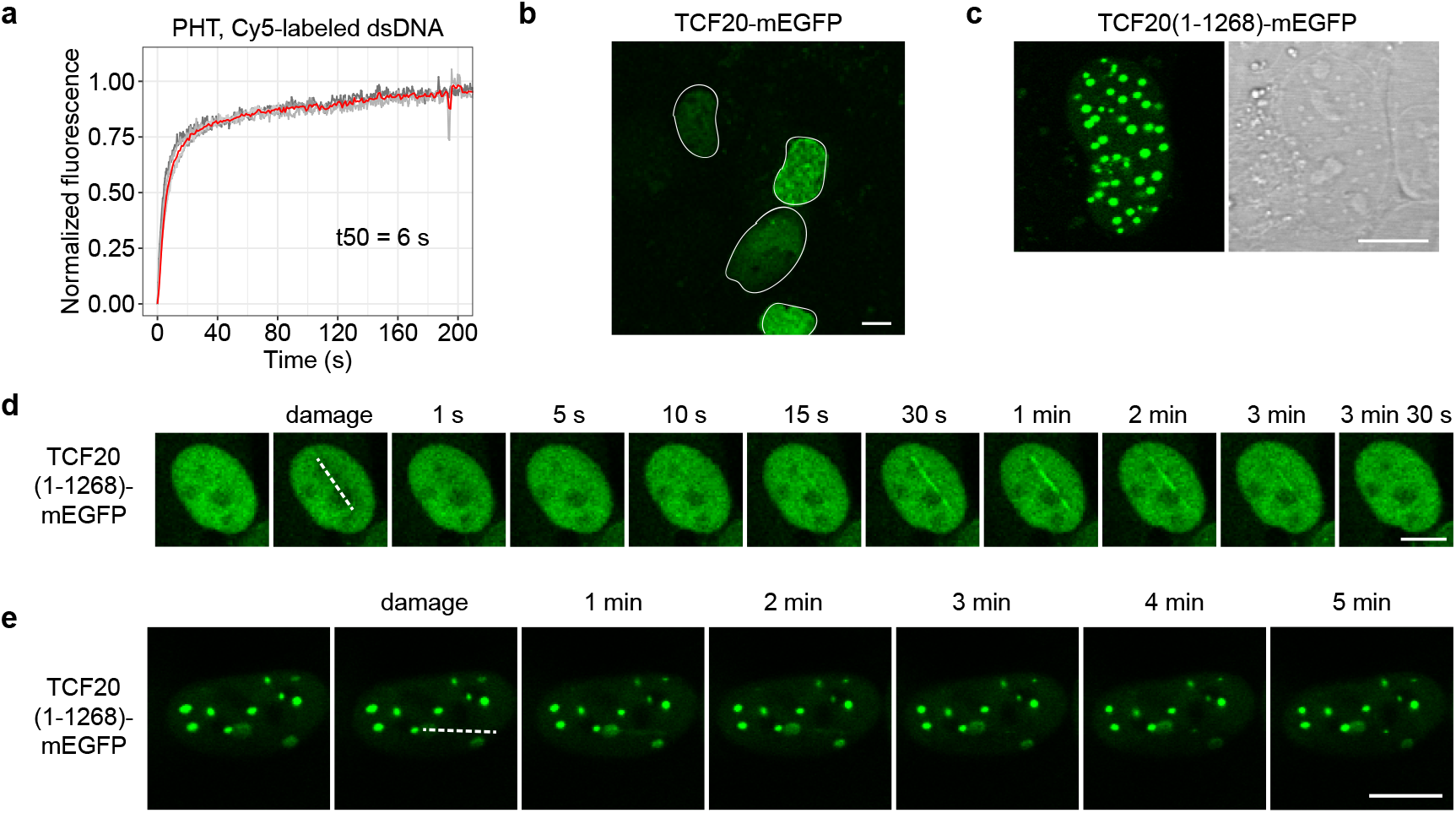
TCF20(1-1268) forms droplets and prematurely dissociates from DNA damage sites. **a,** FRAP kinetics of whole-droplet photobleaching of PHT complex with Cy5-labeled DNA, related to Fig. 7b. Fluorescence intensity right after bleaching is set to 0 and recovery plateau to 1. First-order exponential equation f(t) = 1-exp(−t / τ) did not fit these data. The red line indicates average value of 5 separate curves, binned to 1 s. t50 of this average curve is approximately 6 s. **b**, Full length TCF20-mEGFP transfected into U2OS cells. White outlines indicate nuclear borders. **c**, Confocal microscopy images in GFP and transmission of a U2OS cell transfected with TCF20(1-1268)-mEGFP. **d,** and **e**, Confocal microscope images of TCF20(1-1268)-mEGFP accumulation at DNA damage sites after damage induction with 355 nm laser. **d**: low and **e**: high expression. All scale bars = 10 μm.

**Supplementary Table 1.**
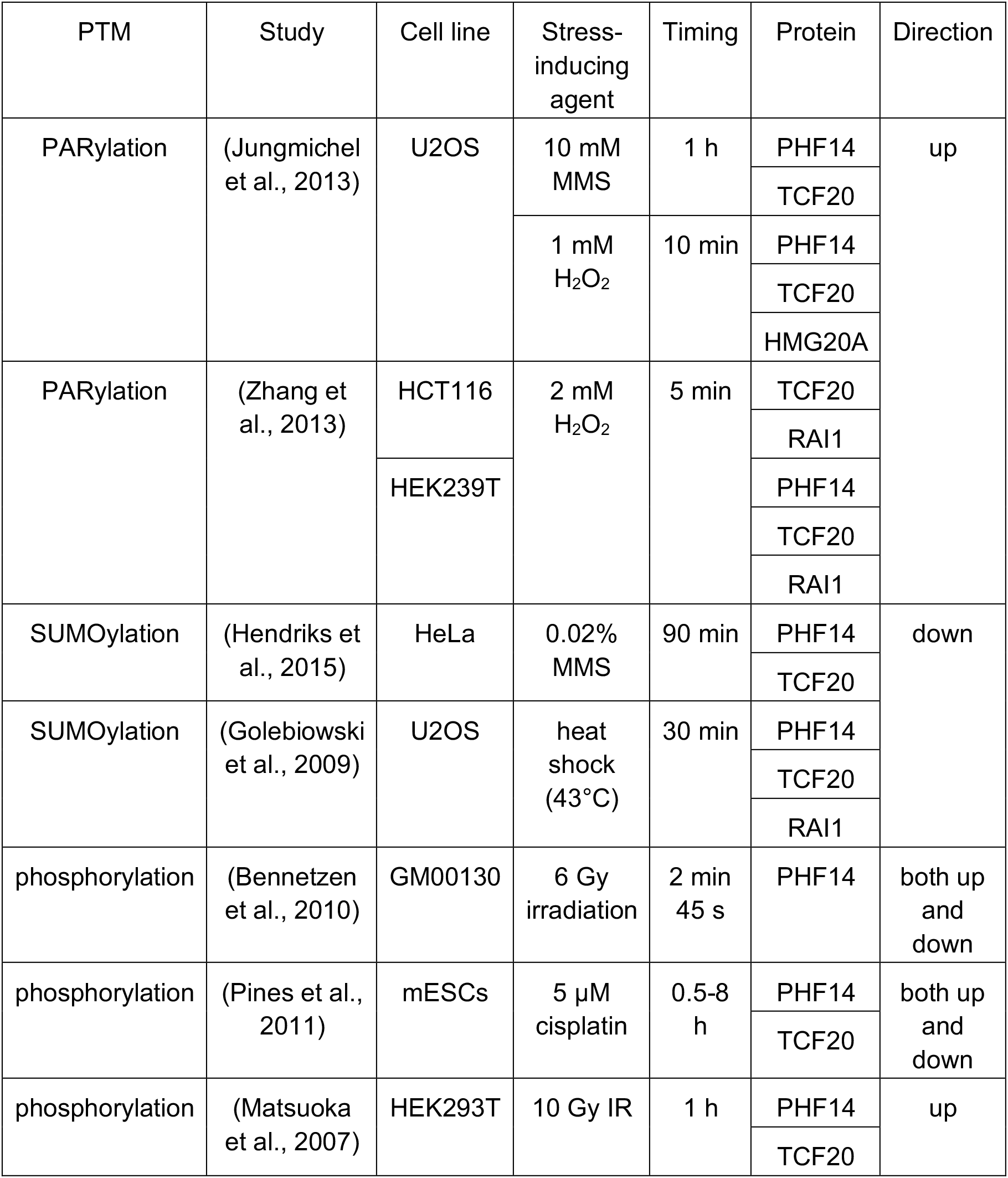
Changes in PTM levels of the PHF14 complex in response to genotoxic stress inducing treatments. Compiled from published literature.

**Supplementary Table 2.**
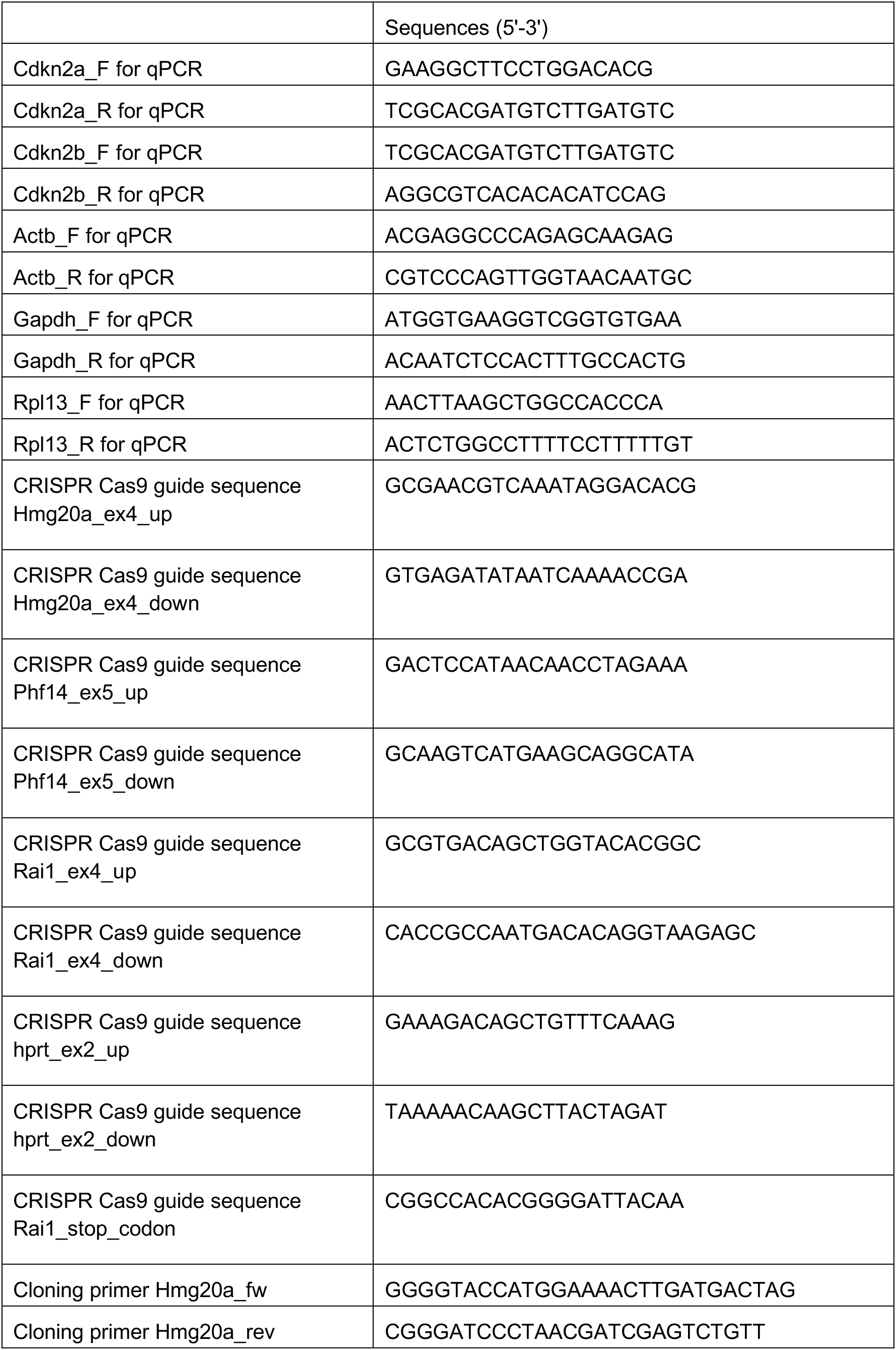
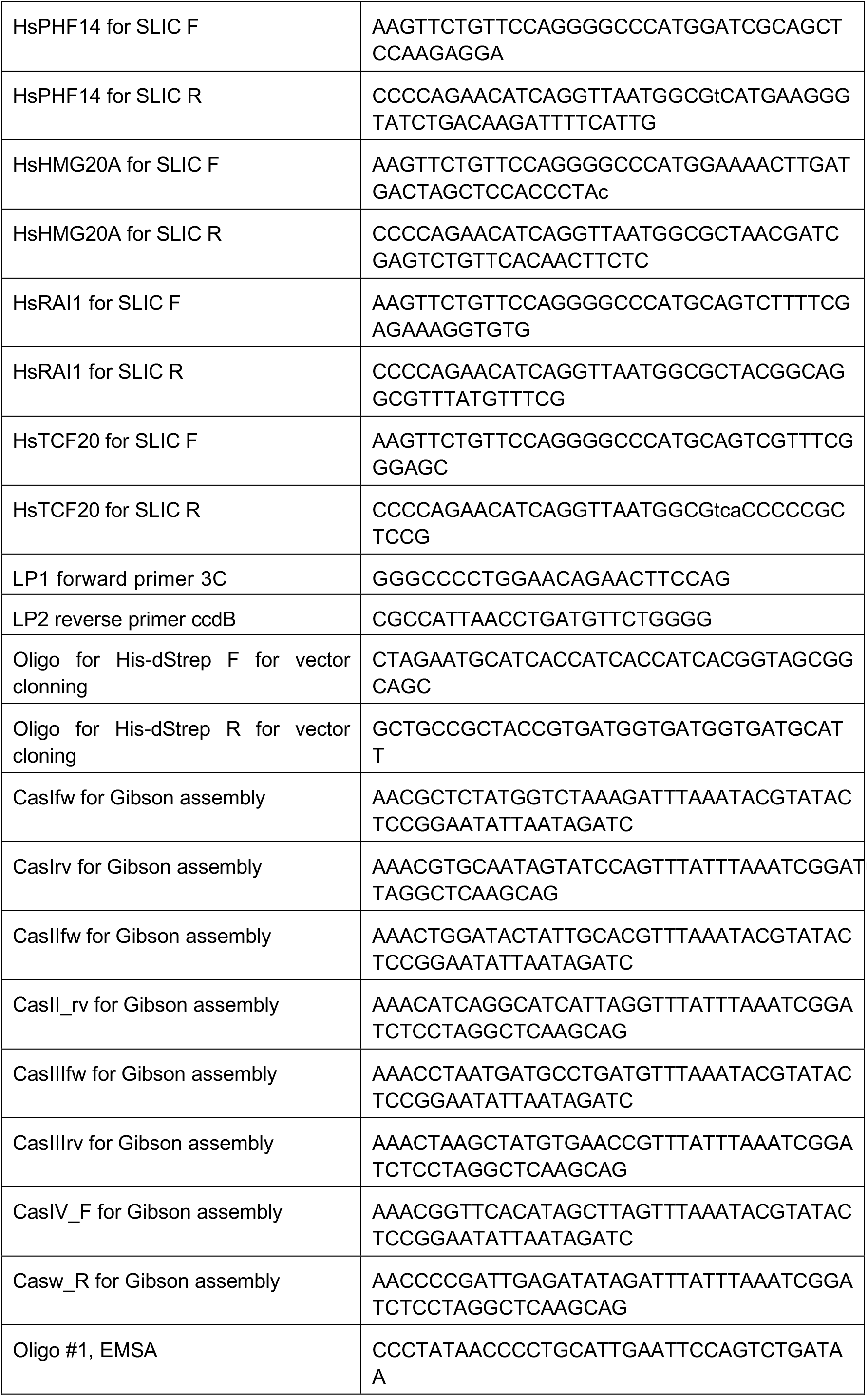
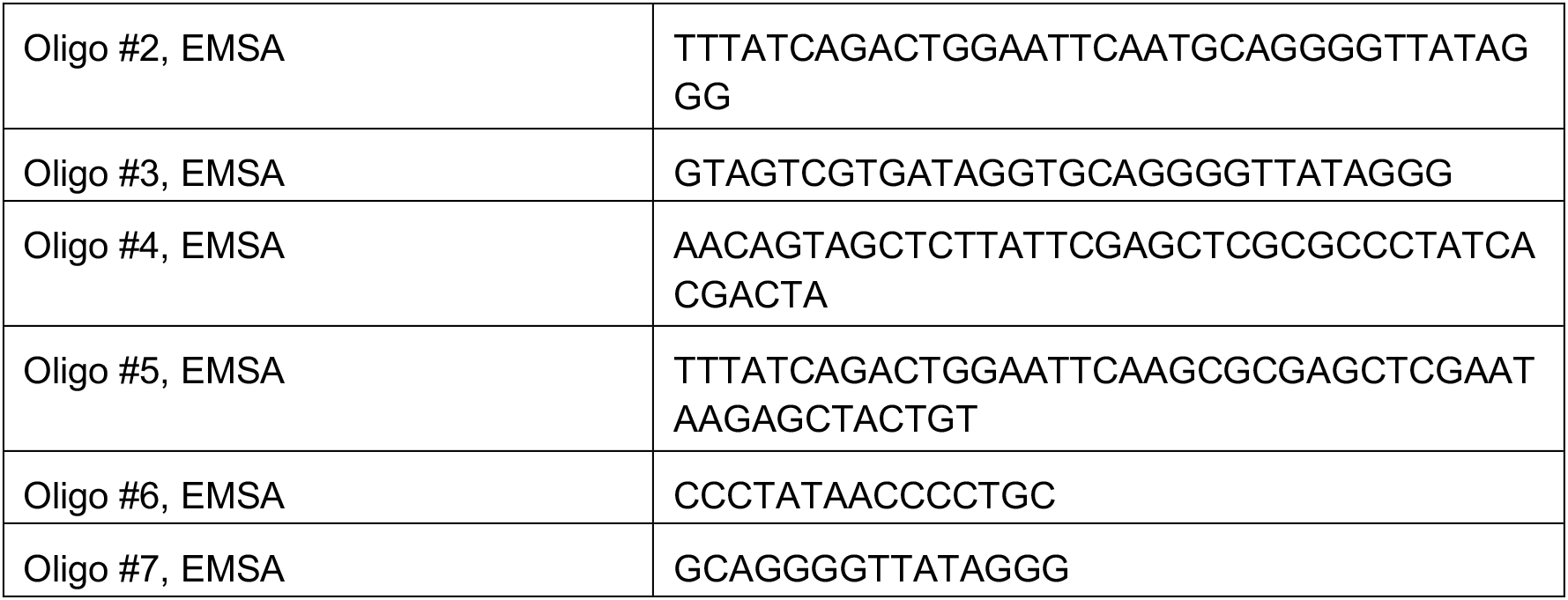
Oligonucleotide sequences used in this study

## Methods

### Cell culture and differentiation

Mouse ESCs (129 × C57BL/6J or clonal lines derived from this background) were cultured in ESC media composed of KnockOut™ DMEM (Gibco), 15% EmbryoMax FBS (Merck Millipore), 1% non-essential amino acids (Gibco), 1% GlutaMAX (Gibco), 1% penicillin/streptomycin (Gibco) and 0.1% β-mercaptoethanol (Sigma-Aldrich) and supplemented with 20 ng/mL leukemia inhibitory factor (Protein Expression and Purification Core Facility, EMBL). Cryopreserved cells were initially plated onto a layer of mouse embryonic fibroblasts (MEFs) and subsequently passaged onto feeder-free, gelatin-coated plates or flasks. Media was changed every day and cells passaged every other day.

Neurons were differentiated from ESCs as previously described^21^. Briefly, ESCs were cultured in low-adhesion Petri dishes (Greiner) in CA media (DMEM high glucose supplemented with 10% FBS, 1% non-essential amino acids, 1% penicillin/streptomycin, 1% GlutaMAX, 1% Na-

Pyruvate and 0.1% β-mercaptoethanol) for 8 days with media changes on days 2, 4 and 6. On days 4 and 6, 5 µM retinoic acid (Sigma-Aldrich) was also added to the media. On day 8, NPCs were collected, dissociated and plated onto poly-D-lysine (Sigma-Aldrich) treated and laminin-coated (Roche) Nunc plates in N2 media composed of DMEM high glucose, 1% Pen/Strep, 1% N-2 Supplement (Thermo Fisher), and 2% B-27™ Supplement (minus vitamin A, Thermo Fisher), which was changed 2 h, 24 h, and 72 h after plating.

For monolayer differentiation of ESCs, cells were plated onto adherent gelatinized plates in CA media with 5 µM retinoic acid at a density of 1-2×105 cells per well of a 6-well plate. Media were changed every two days and cells were considered NPCs from day 4 onwards (assayed by RT-qPCR measurements for increased *Nes*, and decreased *Sox2*, *Pou5f1* and *Nanog* levels).

U-2 OS and HEK293T cells were cultured on adherent plates in DMEM high glucose (Gibco) supplemented with 10% fetal bovine serum (Gibco), 1% penicillin/streptomycin, 1% GlutaMAX and 1% sodium pyruvate. Cells were passaged at 70-90% confluency.

All cells were maintained at 37°C with 5% CO_2_ and routinely tested for *Mycoplasma*. TrypLE Express (Gibco) was used for passaging and harvesting cells. Cell counting experiments were performed using Cellometer Auto T4 (Nexcelom). Live cell percentages were determined using trypan blue staining.

### CRISPR editing

Guide sequences were designed using Benchling or the guide design tool from the Zhang lab (crispr.mit.edu) and cloned into pSpCas9(BB)-2A-GFP (PX458), Addgene #48138 from Feng Zhang, or pSpCas9(BB)-2A-RFP (modified from Addgene plasmid #48138) as described previously^67^. 2×10^6^ ESCs were nucleofected with Cas9-GFP plasmids and a repair template where applicable using 4D-Nucleofector Core Unit (Lonza), following manufacturer’s instructions. Single-cell sorting into MEF-covered 96-well plates using BD AriaFusion was performed 48 hours later to select the cells expressing the fluorescent markers. Surviving cells were expanded for genotyping and freezing. Genomic DNA was extracted either using DirectPCR Lysis Reagent (Viagen Biotech, 301-C) or Puregene Core Kit B (Qiagen). Homozygously edited cells were identified using PCR and Sanger sequencing of gel extracted fragments (gel extraction performed using QIAquick Gel Extraction Kit from Qiagen, according to manufacturer’s instructions) and further verified by western blotting, subject to antibody availability.

The Rai1-HA-3x-FLAG-P2A-GFP cell line was generated by modifying a published method: a donor plasmid was constructed using the pFETCH Donor plasmid (Addgene plasmid #63934 from Eric Mendenhall and Richard M. Myers)^68^ that had been modified to change the selection marker to emGFP and add a HA tag to the protein. The homology arms were inserted using Gibson assembly and guide was designed to target close to the endogenous *Rai1* stop codon. Cells were treated with 10 μM SCR7 (Xcessbio Biosciences) for 24 h before electroporation to increase HDR efficiency.

Phf14 KO (exon 5 deletion), Rai1 exon 4 KO and hprtDRGFP knock-in clonal lines were confirmed to have no chromosomal aberrations either by RNA-seq or DNA-seq^69^. Guide sequences and repair templates are listed in Supplementary Table 2.

### Messenger RNA-sequencing (mRNA-seq)

RNA was extracted from approximately 1×10^6^ cells using the RNeasy Mini kit (Qiagen). DNA was digested using the TURBO DNA-free Kit (Ambion). RNA integrity was assessed using Bioanalyzer (Agilent). Poly(A) selection kit (New England Biolabs) was used to extract mRNA from 1 μg total RNA and NEB Ultra Library Preparation Kit for Illumina was used to prepare sequencing libraries with different indexing primers to enable pooling of up to 10 libraries per lane of 50 bp SE HiSeq 2000 Sequencer (Illumina) runs. Sequencing reads were demultiplexed and mapped to mm10 reference genome using RNA STAR (Galaxy v2.5.2b-0) and read counts per gene were generated using htseq-count (Galaxy Version 0.9.1). DESeq2 (v1.26.0) was used to analyze data and determine differential gene expression.

### MODified histone peptide array

Mouse Rai1 cDNA corresponding to amino acids S1649-L1889 was cloned into pETM11-His-SUMO3 and expressed in BL21-CodonPlus(DE3)-RIL bacteria grown overnight at 16°C in LB with 0.2 mM IPTG and 100 µM zinc-acetate. Cells were harvested by centrifugation, treated with DNAseI and sonicated on ice. Protein was purified using Ni-NTA beads.

ActiveMotif MODified™ Histone Peptide Array (catalog number 13005) was performed according to manufacturer’s recommendations. Briefly, the array was incubated overnight at 4°C in 5% milk in TBS-T, then washed 3x with TBS-T. Purified proteins were added to protein binding buffer (100 mM KCl, 20 mM Tris pH 7.5, 1 mM EDTA, 0.1 mM DTT, 10% glycerol) and the array was incubated with the proteins for 4 h at 4°C. After washing three times, primary antibody solution (Sumo 2 + Sumo 3, abcam, ab3742) or already HRP-conjugated antibody (anti-GST-HRP, Merck RPN1236) was added for 1 h at room temperature, then washed 3x with TBS-T and twice with water. When needed, secondary antibody incubation was performed as well. Immobilon Western Chemiluminescent HRP Substrate was used for imaging with the ChemiDoc Touch imaging system (Bio-Rad, 1708370).

### Protein extraction and western blotting

Crude nuclear lysis was performed to check Phf14 and Hmg20a levels in KO cells. Briefly, 1×10^6^ cell pellets were resuspended in 600 μL 0.5% Triton X-100 and 1x cOmplete protease inhibitor cocktail in PBS and rotated at 10 rpm at 4°C for 30 min. Then, nuclei were pelleted at 5000 rcf for 5 min at 4°C, supernatant was removed, the pellet resuspended in 147 μL of 2x SDS loading dye and DNA was sheared by sonication using the EpiShear Probe Sonicator (Active Motif). 3 μL β-mercaptoethanol was added and samples were heated to 95°C for 10 min before the same volume of each sample was loaded on NuPAGE Novex 4-12% Bis-Tris protein gels and run at 150 V. Precision Plus Protein Western C Standards (Bio-Rad, 161-0376) were used to determine molecular weights. PVDF membranes were used for protein transfer with Mini Blot Modules (Invitrogen) and 10% methanol in the transfer buffer and 5% milk in TBS-T for blocking (1 h at RT) and antibody dilutions (primary antibody incubation overnight at 4°C). 1:3000 or 1:10000 (for H3) dilutions of secondary antibodies (BioRad) (1 h at RT) and Immobilon Western Chemiluminescent HRP Substrate was used for imaging with the ChemiDoc Touch imaging system (Bio-Rad, 1708370).

### Antibodies used for western blotting

| Target | Company | Catalog number | Dilution |
| --- | --- | --- | --- |
| PHF14 | proteintech | 24787-1-AP | 1:1000 |
| HMG20A | proteintech | 12085-2-AP | 1:500-1:1000 |
| HA-Tag | CST | 3724 | 1:1000 |
| Myc-Tag | ThermoFisher | R951-25 | 1:1000 |
| FLAG-Tag | Sigma-Aldrich | SAB4200119 | 1:1000 |
| H3 | abcam | ab1791 | 1:5000 |
| Goat Anti-Rabbit HRP | Bio-Rad | 1706515 | 1:3000-1:10000 |

### Rai1 double affinity purification

Consecutive FLAG and HA pull-downs were performed as described previously^70^. Nuclei were prepared from 2×10^9^ cells by hypotonic lysis and nuclear proteins extracted with 500 mM KCl. The first immunoprecipitation was carried out at 150 mM KCl with 300 μL anti-FLAG resin (Sigma A2220) for 4 h at 4 °C. The resin was transferred to a disposable column, washed with 30 volumes of wash buffer (20 mM HEPES-HCl pH 7.9, 750 mM NaCl, 0.2mM EDTA, 5 mM 2-mercaptoethanol, 10% glycerol, 0.01% NP-40, 0.4 mM PMSF) by gravity flow. Protein complexes were eluted in FLAG-elution buffer (0.25 mg⁄mL FLAG3 peptide, 20 mM HEPES-HCl pH 7.9, 150 mM NaCl, 0.2 mM EDTA, 5 mM 2-mercaptoethanol, 10% glycerol, 0.01% NP-40, 0.4 mM PMSF) and incubated with 50 μL goat anti-HA agarose (Bethyl 190–107) for at least 4 h at 4 °C. The immunoprecipitated material was washed with 10 volumes wash buffer (1 M NaCl; 0.25% NP-40; 0.5 mM DTT; 20 mM HEPES HCl pH 7.9) and eluted 2 times 5 min in 25 μL 20 mM glycine pH 2, 150 mM NaCl. The eluates were submitted for MS analysis at Proteomics Core Facility at EMBL.

### Immunoprecipitation with RAI1 ePHD and PHF14 constructs

20 cm plates of HEK293T cells were transfected with 10 µg of each plasmid using CalPhos transfection kit (Takara) according to manufacturer’s instructions and harvested 3 days later. Pellets were resuspended in 600 μL hypotonic buffer (20 mM HEPES-HCl pH 7.9, 10 mM NaCl, 1 mM EDTA, 10% glycerol, 0.1% NP-40, 0.4 mM PMSF, 1 mM DTT supplemented with cOmplete protease inhibitor cocktail), rotated at 4°C for 10 min. NaCl was added to bring the concentration to 150 mM. Lysates were sonicated 30 s on / 30 s off for 10 min in a bath sonicator, then centrifuged and supernatant collected. The remaining pellet was resuspended in 300 μL IP buffer (20 mM HEPES-HCl pH 7.9, 150 mM NaCl, 1 mM EDTA, 10% glycerol, 0.4 mM PMSF, 1 mM DTT, 1x cOmplete) and centrifuged. The supernatant was added to the previously collected and more IP buffer was added to dilute NP-40.

Myc-Tag (9B11) antibody (NEB 2276 S) and mouse IgG (Santa Cruz sc-2025) were coupled to Protein G Dynabeads (Invitrogen 10004D) according to manufacturer’s instructions or FLAG M2 beads (Sigma Aldrich) beads were used. Lysates were added to the beads and left at 4°C on rotation overnight. IP buffer with 200 mM NaCl and 0.01% NP-40 was used to wash the beads four times (including one 10 min wash on rotation). Immunoprecipitated proteins were eluted in 4x SDS sample buffer with 5% β-mercaptoethanol at 95°C for 5-15 min.

### Phf14 immunoprecipitation and liquid chromatography–mass spectrometry

100 million ES cells or NPCs were collected per condition with 2-3 clonal lines of Phf14 KO cells used. Pellets were frozen in liquid nitrogen and stored at −80°C. Thawed pellets were resuspended in 5 mL hypotonic buffer (15 mM HEPES pH 7.9, 5 mM MgCl_2_, 30 mM KCl, 3 mM CaCl_2_, 1 mM DTT, 0.8 mM PMSF, 1 mM benzamidine, 1 mM sodium metabisulfite), followed by 15 strokes with a dounce homogenizer. Cell disruption was checked by trypan blue staining. Nuclei were collected by centrifugation (700 rcf, 10 min) and were resuspended in 2 mL MNase digestion buffer (15 mM HEPES pH 7.9, 5 mM MgCl2, 10 mM KCl, 0.5 mM EDTA, 1 mM DTT, 0.8 mM PMSF, 2x cOmplete). 28 U MNase (Worthington) per 50 million cells was used to fragment DNA at 37°C for 4 min (500 rpm shaking). Reaction was stopped by adding EDTA (5 mM final concentration) and EGTA (3 mM) and Triton X-100 (0.02%) to disrupt nuclear membranes with dounce homogenizer (20 strokes) or pipetting. Supernatant was collected after centrifugation (17,000 × g, 10 min) and glycerol was added to a final concentration of 5%. DNA fragmentation patterns were confirmed to correspond to mostly mono- and oligonucleosomes.

Phf14 antibody (proteintech 24787-1-AP) was coupled to M-280 Sheep Anti-Rabbit IgG Dynabeads (Invitrogen 11203D) according to manufacturer’s instructions overnight (7 µL antibody, 50 µL beads per IP). Immunoprecipitation was performed overnight on rotation. The magnetic beads were then washed 4 times with wash buffer (20 mM HEPES pH 7.9, 200 mM NaCl, 0.01% NP-40, 1 mM EDTA, 10% glycerol, 1 mM DTT, 0.8 mM PMSF, 1x cOmplete) and the proteins were eluted in SDS sample buffer with 2% β-mercaptoethanol at 95°C for 10 min. All steps were performed either at 4°C or on ice, unless otherwise noted.

Proteins were separated on an SDS-PAGE and bands corresponding to immunoglobulins (25 kDa and 50 kDa) were removed and not analyzed by mass spectrometry (this included a 45 kDa band in ESCs that also included some Hmg20a according to a separate MS analysis run). The experiment with etoposide treatment was performed in the same manner. The cells were treated with 5 µM etoposide (Sigma-Aldrich E1383) for 1 h. The eluted proteins in SDS sample buffer were submitted directly for MS analysis without SDS-PAGE.

Samples were prepared at the EMBL-Heidelberg Proteomics Core Facility with the SP3 protocol^71^ and trypsin digested peptides analyzed by liquid chromatography followed by tandem mass tag (TMT)-labeled mass spectrometry using TMT10plex isobaric label reagent set (ThermoFisher). Raw data were mapped and quantified using MaxQuant (v1.6.3.4) (Cox and Mann, 2008). A variance stabilization normalization method was applied with the R package vsn (v3.45)^72^. Limma (v3.42)^73^ and fdrtool (v1.2)^74,75^ were used to analyze data in R.

### Cell cycle analysis

Click-iT EdU Alexa Fluor 488 Flow Cytometry Assay Kit (C10425, Invitrogen) was used for cell cycle analysis, following manufacturer’s instructions. 10 µM EdU was added to the media of monolayer-differentiated NPCs for 2 h before cells were harvested for analysis. DNA content was assayed with 1 μg/mL final concentration of DAPI in 500 μL 1 x Click-iT saponin-based permeabilization and wash buffer. Fluorescence was measured using BD LSRFortessa™ and results were analyzed using FlowJo.

### RT-qPCR

RNA was harvested from cell pellets of 1 million cells using the RNeasy Mini kit (Qiagen). DNA was digested using the TURBO DNA-free Kit (Ambion) and cDNA was synthesized using High-Capacity cDNA Reverse Transcription Kit (Applied Biosystems). Quantitative PCR was performed on a QuantStudio 6 Flex Real-Time PCR System using SYBR Green Power SYBR™ Green PCR-Master-Mix (Applied Biosystems™) and primers listed in Supplementary Table 2.

### DNA fiber spreading

DNA fiber spreading experiments were performed as previously described^34,35^. Briefly, cells were grown in 6-well plates to approximately 50% confluency. 50 μM IdU (Cayman Chemical 20222) was added to the media and cell were incubated at 37°C for 20 min, then for another 20 min in 250 μM CldU (Cayman Chemical 18155). Depending on the assay, 4 h in 1 mM hydroxyurea step was included after those steps. Between each step, cells were washed twice in PBS. Afterwards, cells were trypsinized and resuspended in PBS to a concentration of 150 to 1000 cells/μL. 2 μL of cell suspension was dropped on a microscope slide and allowed to dry for 10 min. Cells were lysed on the slide with 9 uL lysis buffer (0.5% SDS, 200 mM Tris– HCl pH 7.5, 50 mM EDTA) for 10 min and then slides were tilted to allow the DNA to spread down the microscope slide. After air drying, the slides were fixed in 3:1 methanol:acetic acid for 10 min, dried and stored at 4°C. DNA was denatured in 2.5 M HCl and blocked in 5% BSA in PBS. Primary antibodies against IdU and CldU were diluted in 5% BSA at 1:100 (BD Biosciences 347580) and 1:250 (abcam ab6326) and secondary antibodies at 1:1000 (Alexa Fluor 488 anti-mouse, Invitrogen A-11001) or 1:125 (DyLight 594 anti-rat, Invitrogen SA5-10020). Fluoromount-G™ (Invitrogen 00-4958-02) or ProLong Gold (ThermoFisher P10144) were used for mounting coverslips on the slides. Slides were imaged with a 60X oil-immersion objective on a Nikon Ti-E widefield microscope and images were analyzed using Fiji. All non-overlapping tracks with IdU (first label) signal were measured in both channels.

### Comet assay

ESCs were differentiated to neurons in 6-well plates, treated with 5 μM etoposide (Sigma-Aldrich E1383) for 1 h and harvested together with untreated controls by scraping on day 12. Alkaline comet assay was performed following manufacturer’s instructions using the CometAssay Kit (Trevigen, 4250-050-K). Gel electrophoresis was performed at 4°C using Wide Mini-Sub Cell GT Horizontal Electrophoresis System (BioRad, 1640301) at 14 V (approximately 300 mA) for 35 min. SYBR Gold (Invitrogen) was used to visualize DNA. Images were acquired on Nikon Ti-E widefield microscope. CometScore 2.0 software was used to analyze images and results were manually verified to exclude incorrectly marked comets.

### DR-GFP reporter assay and generation of I-SceI lentivirus

DR-GFP cell lines were generated as described under CRISPR editing and in previous reports^36^, using pHPRT-DRGFP (Addgene plasmid #26476 from Maria Jasin) as a repair template and two guides targeting the X chromosome near the homology arms start sites (2×10^6^ ESCs were nucleofected with 1 μg of each guide plasmid and 3 μg of repair template and plated onto 3 wells of DR3 puromycin-resistant MEF in a 6-well plate). Positive integration events were enriched for by 5 days of puromycin selection (1 μg/mL) followed by 5 days of 6-thioguanine selection (10 μg/mL). At that point, colony selection was performed and colonies were trypsinized and moved to a 96-well and expanded to 6-well plates. Correct integration from both ends of the construct was confirmed by PCR amplification of genomic DNA.

I-SceI enzyme was introduced to the cells by lentivirus. Lentivirus was produced in HEK293T cells using a 2^nd^ generation lentiviral packaging system (psPAX2 – Addgene plasmid #12260 and pMD2.G – Addgene #12259 from Didier Trono) and pRRL.EF1a.HA.NLS.Sce(opt).T2A.BFP (Addgene #32628 from Andrew Scharenberg) and CalPhos transfection kit according to manufacturer’s instructions (Takara). 2 days after transfection, HEK293T cells were checked for BFP expression under a microscope with 90% or more expected to be successfully transfected. The medium was collected and centrifuged at 700 rcf for 10 min, then filtered through a 0.45 μm filter to remove large debris and then centrifuged at maximum speed using a SW 28 Ti Swinging-Bucket Aluminum Rotor (Beckman Coulter) for 2-4 h. The viral pellet was then resuspended in 100 μL PBS, aliquoted and stored at −80°C. To determine the amount of virus required for experiments, a titration experiment was performed, using flow cytometry to measure BFP expression. Transduction efficiency in final experiments was between 20-60%.

### Laser microirradiation

For live-cell imaging of fluorescently tagged proteins, Olympus FV1200 microscope was used with 60X water immersion objective (N/A 1.20). 355 nm pulsed laser was used through SIM scanner to generate DNA damage (45-50%, 1 ms/pixel, 1024×1024 pixels, zoom 1). GFP-tagged proteins were imaged with 488 nm argon laser and mRuby/mCherry-tagged proteins with 559 nm laser. Cells were transfected with plasmids encoding fluorescently tagged proteins using FuGENE® HD Transfection Reagent (Promega), following manufacturer’s protocol and assayed 1-3 days after transfection. Cells were grown on ibiTreat 8-well µ-slides or 35 mm µ-dishes (ibidi, 80826 and 81156) that were either untreated or coated with 0.1% gelatin for 15 min for mouse cells. During imaging, cells were kept in a humidified chamber with temperature and CO_2_ regulation (37°C, 5% CO_2_). Data quantification was performed using Fiji and analysis in R.

For immunofluorescence-based DNA damage analysis, LSM780 NLO was used with water immersion objective “C-Achroplan” 32x/0.85 W Corr M27. Every 10^th^ line of 124×124 pixel areas (zoom 1) was scanned with 800 nm 2-photon laser (7-10% laser power) to generate damage. Cells were subsequently fixed and analyzed by immunofluorescence staining using Nikon Ti Eclipse widefield microscope.

### Immunofluorescence

Cells were grown on ibiTreat 8-well µ-slides or 35 mm µ-dishes (ibidi, 80826 and 81156) that were either untreated or coated with 0.1% gelatin for at least 15 min for mouse cells and fixed in 3% PFA for 20 min at room temperature, PFA was then quenched for 5 min in 30 mM glycine. Cell were then washed twice in PBS, permeabilized with 0.1% Triton-X 100 for 10 min, washed again in PBS and blocked for 30 min in 0.5% BSA. Primary antibody (manufacturers’ recommended dilutions from datasheets in 0.5% BSA, if a range or none provided, 1:200 dilution was used) incubation was performed overnight at 4°C on a shaker. Next day cells were washed three times in PBS, then incubated with secondary antibodies at recommended dilutions for 30-60 min at room temperature with shaking, washed again twice with PBS, incubated with DAPI (5 µg/mL in PBS) for 5 min, then washed again in PBS and left in PBS for imaging. Samples were imaged with Zeiss LSM780 (NLO) confocal microscope or Nikon Ti-E widefield microscope.

Cells were grown on ibiTreat 8-well µ-slides or 35 mm µ-dishes (ibidi, 80826 and 81156) that were either untreated or coated with 0.1% gelatin for at least 15 min for mouse cells and fixed in 3% PFA for 20 min at room temperature, PFA was then quenched for 5 min in 30 mM glycine. Cell were then washed twice in PBS, permeabilized with 0.1% Triton-X 100 for 10 min, washed again in PBS and blocked for 30 min in 0.5% BSA. Alternatively, fixation and permeabilization with methanol at −20°C for 4 min was used. Primary antibody (manufacturers’ recommended dilutions from datasheets in 0.5% BSA, if a range or none provided, 1:200 dilution was used) incubation was performed overnight at 4°C on a shaker. Next day cells were washed three times in PBS, then incubated with secondary antibodies at recommended dilutions for 30-60 min at room temperature with shaking, washed again twice with PBS, incubated with DAPI (5 µg/mL in PBS) for 5 min, then washed again in PBS and left in PBS for imaging. Samples were imaged with Zeiss LSM780 (NLO) confocal microscope or Nikon Ti-E widefield microscope.

### Antibodies used for immunofluorescence

| Target | Company | Catalog number | Dilution |
| --- | --- | --- | --- |
| PHF14 | proteintech | 24787-1-AP | 1:200 |
| HMG20A | proteintech | 12085-2-AP | 1:200 |
| HA-Tag | CST | 3724 | 1:800 |
| phospho-histone H2A.X | Merck | 05-636 | 1:1000 |
| phospho-histone H2A.X | CST | 9718 | 1:400 |

### FRAP

FRAP experiments were performed similarly to laser microirradiation using Olympus FV1200 microscope. For photobleaching, 488 nm laser was used at maximum power for 1-3 frames, as necessary. Data quantification was performed using Fiji and analysis in R.

### Molecular cloning and plasmids

PHF14, RAI1 and TCF20 cDNA sequences were obtained from Promega (products FXC01995, FHC01638, FXC11595). RAI1 ePHD domain was codon-optimized for bacterial expression. HMG20A cDNA sequence was obtained by PCR from cDNA generated from RNA extracted from HeLa cells using the RNeasy Mini kit (Qiagen) and ProtoScript II First Strand cDNA Synthesis Kit (NEB) with Oligo-dT primers. Q5 Site-Directed Mutagenesis Kit (NEB E0554S) and restriction cloning were used to generate partial constructs. mRuby was amplified from PB-EF1α-MCS-IRES-RFP (System Biosciences PB531A-2) by PCR. PCR amplified sequences were inserted into pcDNA3.1(+) (Invitrogen), pcDNA3 (Invitrogen), pmEGFP-N1/pmEGFP-C1 (kind gifts from Jennifer Lippincott-Schwartz) or pmCherry-N1 (Clontech 632523). GFP-ZMYND8 (Addgene 65401)^40^ and EGFP-N1-PARP1 (kind gift from Andreas Ladurner) were used as positive controls for laser microirradiation experiments. Relevant primers sequences are listed in Supplementary Table 2.

### TCF20(1-1268) construct experiments

TCF20(1-1268)-mEGFP construct was generated from the TCF20-mEGFP plasmid using the Q5 Side-Directed Mutagenesis Kit (NEB, E0554S) following manufacturer’s instructions. The cells were imaged live using Olympus FV1200 microscope with 60X water-immersion objective (NA 1.20). For 1,6-hexanediol treatments, 1,6-hexanediol (Sigma 240117) was diluted in DMEM to achieve 10% concentration (w/v) with 10 µg/ml digitonin (Merck D141). Equal volume of the HD solution was added to the cell media while imaging, yielding a final concentration of 5% HD and 5 µg/ml digitonin.

### DNA constructs for recombinant protein expression in insect cells

ORFs of human PHF14, HMG20A, RAI1 and TCF20 (see Molecular cloning and plasmids section for details) were cloned into pCoofy vectors using Sequence and Ligation Independent Cloning (SLIC)^76,77^. pCoofy27 (N-His7) was provided by the Protein Expression and Purification Core (Pepcore) facility in EMBL. N-His6-double Strep-pCoofy vector was generated from pCoofy51 (N-dStrep) by insertion of a His6-coding sequence into XbaI and Eco47III sites of pCoofy51.

Each ORF and pCoofy vector were linearized with gene-specific SLIC primers and LP1/LP2, respectively, by PCR. Amplified DNA was extracted using QIAquick Gel Extraction Kit and mixed 1:1 ratio of vector and insert for SLIC reaction.

For multigene expression, gene cassettes encoding N-His6-dStrep-PHF14, and N-His7-HMG20A, N-His7-RAI1 and N-His7-TCF20 constructs were co-inserted into the pBig1a vector by Gibson assembly as described^78^. Assembled DNAs were transformed into Top10 chemically competent cells. The insertion of each DNA was confirmed by Swa1 restriction enzyme digestion and Sanger sequencing.

pCoofys and pBig1a vectors harboring genes for insect cell proteins expression for next steps were subjected to bacmid transformation using DH10EMBacY E. coli competent cells (Geneva Biotech) with electroporation. Only white colonies were selected from Tet/Kan/Gent/Amp, IPTG, XGal plates. Bacmids were extracted with isopropanol precipitation. All the primer sequences used in this section are listed in Supplementary Table 2.

### Insect cell culture and baculovirus generation

Sf21 cells were grown in Sf-900 III Serum-Free Media (Gibco, 12659017) in insect cell culture room at 30 °C in atmosphere. Cells were passaged for every 2-3 days to 0.3-0.5 x 10^6^ /mL cells. 10 μg of extracted bacmid were transfected into 0.9 x 10^6^ of Sf21 cells using X-tremeGENE HP DNA Transfection Reagent. After 3 days, the supernatants were collected and used to generate V1. At a viability of around 85% the supernatant was collected again, and used to infect Sf21 cells. Optimized infection ratios were selected for each batch of baculoviruses to have viability of 90% after 2-3 passages (1.3-1.8 x 10^6^ /mL). Cells were harvested at 600 g and the pellets were stored in −80 °C.

### Protein purification

Proteins were purified from Sf21 insect cells by affinity purification with Ni-NTA agarose (Qiagen, 30230) and, where possible, Strep-Tactin sepharose (Iba, 2-1201) beads, followed by ion exchange chromatography and size exclusion chromatography on an ÄKTA Pure system (GE Healthcare). Sf21 cell pellets were resuspended in lysis buffer, 50 mM Tris-HCl (pH 7.5), 500 mM NaCl, 30 mM imidazole, 2 mM β-mercaptoethanol, 4 mM MgCl_2_, 10% glycerol with protease inhibitor cocktail (Sigma, S8830) and Benzonase (Millipore, 70746-3). The lysates were homogenized by sonication and centrifuged for 1 h, 45.000 rpm in an ultracentrifuge (Beckman). For purification of PHF14/HMG20A, cells overexpressing each protein were re-suspended together for co-lysis. The cleared lysates were incubated with pre-equilibrated Ni-NTA agarose beads for 1h and washed twice with washing buffer, 50 mM Tris-HCl (pH 7.5), 500 mM NaCl, 30 mM imidazole and 2 mM β-mercaptoethanol, once with high salt washing buffer, 1 M NaCl, 30 mM imidazole and 2 mM β-mercaptoethanol, and again twice with washing buffer. Elution was performed with 300 mM imidazole, 50 mM Tris-HCl (pH 7.5), 200 mM NaCl and 2 mM β-mercaptoethanol. Where a Strep-tag was present, eluate was incubated with Strep-Tactin sephprose beads for 1 h, washed 3 times with washing buffer, 50 mM Tris-HCl (pH 7.5), 200 mM NaCl, 2 mM β-mercaptoethanol, 0.05 μg/mL of avidin (IBA, 2-0204-015), and eluted with 10 mM biotin. Protein concentration was measured and digested with His-3C protease (Pepcore, EMBL), 1:100 (w/w), overnight at 4 °C.

Subsequently, PHF14, PHF14/HMG20A and ternary and quaternary PHF14 complexes were applied to 5 mL HiTrap Q HP columns. HMG20A was applied to a HiTrap SP HP columns (GE Healthcare GE29-0513-25 and GE29-0513-24) for ion exchange chromatography. Proteins were eluted over a 200 mM – 2 M NaCl gradient in 20 mM Tris-HCl, pH 7.5, 2 mM DTT. Eluted peak fractions were loaded onto SDS-PAGE to check the proteins and concentrated with Amicon Ultra-4 centrifuge filter (Millipore, UFC3080) and used for next step. Size exclusion chromatography was done in 20 mM Tris-HCl (pH 7.5), 200 mM NaCl and 2 mM DTT on a Superdex 200 column (GE Healthcare). In case of individual purification of PHF14 and HMG20A, 300 mM of L-Arginine-HCl was added to the running buffer. Eluted proteins were analyzed on SDS-PAGE and concentrated with Amicon Ultra concentrators (Millipore), aliquoted and stored at −80 °C. Optimized conditions for each protein are described below when needed. All purified proteins were confirmed for their identifications by LC-MS/MS.

### Sample preparation for cross-linking mass spectrometry

Protocols are adopted from Proteomic Core Facility (PCF) in EMBL. 50 ug of purified protein complex at1 ug/ul concentration in 50 mM HEPES (pH 8.5) buffer were incubated with 1mM of DSS-H12/D12 (Creative Molecules Inc., 001S) to have 1 mM in final for 30 min at 35°C, 600 rpm. Cross linking was quenched by adding 0.1 volume of 1 M of ammonium bicarbonate (AmBic) for 10 min at 35°C 600 rpm. To prepare conditions for LysC digestion, 0.8 volume of 10 M Urea, 250 mM Ambic and 0.05 volume of resuspended RapiGest with 10 mM of Ambic were added. The solutions were sonicated for 1 min, DTT was added to have 10 mM in final and then incubated for 10 min at 37°C, 600 rpm. 100 mM of Iodoacetamide was added to have 15 mM and incubated for 30 min at RT in the dark. For digestion, 0.1 μg/μl of Lysyl Endopeptidase (Wako, WDH 4348) in 10 mM of Ambic was used in 1:100 ratio (Protease : Protein) and incubated for 3-4 h at 37°C 600 rpm. By adding HPLC-H_2_O, the urea was adjusted to have 1.5 M and then trypsin (Promega, V511A, sequencing grade) was added in 1:50 ratio (Protease : Proteins) and incubated at 37 °C for 4 h for overnight. Digestion was stopped by adding TFA to 1% in final and incubation for 30 min. at 37°C. The lysates were cleared by centrifugation at 17.000 g at 4°C and transferred to a new tube. Samples were further analyzed at the PCF, EMBL. Pepcore for further analysis. XL-MS data were visualized with xiNET^79^ (open source, http://crosslinkviewer.org/).

### Liquid chromatography-mass spectrometry/mass spectrometry (LC-MS/MS)

Sample preparation and Liquid chromatography-mass spectrometry/mass spectrometry (LC-MS/MS) were performed by PCF in EMBL. Briefly, cysteines were reduced using 10 mM of DTT at 56 °C for 30 minutes and then cooled to 24 °C. Alkylation was done with 20 mM of 2-chloroacetamide RT, in the dark, 30 min. Samples were prepared for the LC-MS/MS with SP3 protocol^71^ and digested with trypsin. Cleaning up of the peptides were performed using OASIS HLB µElution Plate (Waters).

An UltiMate 3000 RSLC nano LC system (Dionex), a trapping cartridge (µ-Precolumn C18 PepMap 100, 5µm, 300 µm i.d. x 5 mm, 100 Å) and an analytical column (nanoEase™ M/Z HSS T3 column 75 µm x 250 mm C18, 1.8 µm, 100 Å, Waters) coupled to Orbitrap Fusion Lumos (Thermo) mass spectrometer were used for peptide separation and ionization (positive ion mode).

The resolution of MS scans was 120,000 and the filling time was a maximum of 50 ms. The resolution of MS2 scans was 15,000 with filling time of 54 ms and a target of 2×10^5^ ions. Data was acquired with data dependent mode. HCD normalized collision energy was set to 34.

### Data processing for LC-MS/MS

Data processing was performed by PCF in EMBL. The raw mass spectrometry data was processed with MaxQuant (v1.6.3.4) (Cox and Mann, 2008). The mass error tolerance was set to 20 ppm and for the MS/MS spectra to 0.5 Da. A maximum of 2 missed cleavages was allowed. For protein identification a minimum of 2 unique peptides with at least seven amino acids and a false discovery rate below 0.01 were required. Quantification was performed using iBAQ values^80^ from the protein lysates. The values are calculated as the sum of the intensities of the identified peptides.

### Phase separation assays

For dilution of proteins, 50 mM Tris-HCl (pH 7.5) and 1 mM DTT with 0 to 500 mM of NaCl buffers were used and polyethylene glycol (PEG) 8000 was added to a final concentration of 5-10%. For imaging chambers, microscopic glass slides and coverslips were washed 1x with water, 1x with ethanol and 1x with water and 2 µl of protein solution was placed between the glass and coverslips using double-sided tape right after mixing ^50^. For measuring turbidity, clear 384-well plates were used and OD was measured at 600 nm with Tecan Fluorescence Microplate Reader.

### Electrophoretic Mobility Shift Assay (EMSA)

The oligomer sequences used for EMSA assays are listed in Supplementary Table 2^45^. The oligos were annealed to have 10 µM final concentration using a thermocycler first by dissolution at 95 °C for 5 min and decrementing the temperature gradually to RT, stored at − 20°C and diluted to desired final concentration. Annealing buffer contained 10 mM Tris-HCl, pH 7.5, 50 mM NaCl and 1 mM EDTA. Non-denaturing gels of 4% acrylamide were made with 0.4 M Tris-HCl, pH 8.45, sucrose 438 mM and 10% APS and TEMED. The migration buffer contained 27 mM Tris and 191 mM glycine. The gels were pre-ran for 1 h at 100 V before running. Proteins were visualized with Coomassie blue and DNA with SYBR Gold.

### Negative stain

Purified proteins (0.01 mg/mL), diluted in 50 mM Tris-Cl, pH 7.5, 150 mM NaCl, 1 mM DTT, was deposited on carbon-coated grids (Electron Microscopy Sciences, 215-412-8400) and stained with 2% (w/v) uranyl acetate. Data were collected at 120 kV with a pixel size of 6.876 Å/px, defocus range of −1.5 to −2.5 μm and total electron dose of 20 e-/Å2. Micrographs for negative stain were obtained with a Tecnai T12 equipped with a 4K CCD camera and Serial EM.

## Notes

### Competing Interest Statement

The authors have declared no competing interest.

